# Knockdown proteomics reveals USP7 as a regulator of cell-cell adhesion in colorectal cancer via AJUBA

**DOI:** 10.1101/2024.10.11.617864

**Authors:** Ahood Al-Eidan, Ben Draper, Siyuan Wang, Brandon Coke, Paul Skipp, Yihua Wang, Rob M. Ewing

**Affiliations:** School of Biological Sciences, Faculty of Environmental and Life Sciences, University of Southampton, Southampton, United Kingdom; Department of Biology, College of Sciences, Imam Abdulrahman Bin Faisal University, P.O. Box 1982, Dammam, Saudi Arabia

## Abstract

Ubiquitin-specific protease 7 (USP7) is implicated in many cancers including colorectal cancer in which it regulates cellular pathways such as Wnt signalling and the P53-MDM2 pathway. With the discovery of small-molecule inhibitors, USP7 has also become a promising target for cancer therapy, and therefore systematically identifying USP7 deubiquitinase interaction partners and substrates has become an important goal. In this study, we selected a colorectal cancer cell model that is highly dependent on USP7 and in which USP7 knockdown significantly inhibited colorectal cancer cell viability, colony formation, and cell-cell adhesion. We then used inducible knockdown of USP7 followed by LC-MS/MS to quantify USP7 dependent proteins. We identified the Ajuba LIM domain protein as an interacting partner of USP7 through co-IP, its substantially reduced protein levels in response to USP7 knockdown, and its sensitivity to the specific USP7 inhibitor FT671. The Ajuba protein has been shown to have oncogenic functions in colorectal and other tumours, including regulation of cell-cell adhesion. We show that both knockdown of USP7 or Ajuba results in a substantial reduction of cell-cell adhesion, with concomitant effects on other proteins associated with adherens junctions. Our findings underlie the role of USP7 in colorectal cancer through its protein interaction networks and show that the Ajuba protein is a component of USP7 protein networks present in colorectal cancer.

## INTRODUCTION

There has been rapidly increasing interest in the role of deubiquitinating enzymes (DUBs) in many types of tumour including colorectal cancer (CRC) (Dewson et al., 2023). DUB proteins are proteases that reverse E3 ligase activity by cleaving a single ubiquitin, polyubiquitin chains or ubiquitin-like modifications on target proteins (Bushman et al., 2021; Reyes-Turcu et al., 2009; Sowa et al., 2009). This role of DUBs is similar to the regulatory role of the phosphatases in a kinase/phosphatase pathway(Reyes-Turcu et al., 2009). There are nearly 100 DUBs in the human genome, grouped into five families: USP, OUT, UCH, Josephin (cysteine proteases) and JAMM (a family of metalloproteases); DUBs have high specificity for ubiquitin. Different DUBs have different domains for the protein-protein interactions, substrate specificity and cellular localisation. Quantitative studies of multiple aspects of DUBs, including protein activity, protein network and genetic studies, have discovered new DUB functions in both humans and yeast (Reyes-Turcu et al., 2009)

Deubiquitination is a regulatory process involved in multiple cell functions, such as cell cycle regulation (Song and Rape, 2008) gene expression (Daniel and Grant, 2007) and DNA repair (Kennedy and D’Andrea, 2005). Moreover, mutations in many DUBs proteins have been implicated in various types of disease, including cancer and neurological disorders (Fischer, 2003; Shanmugham and Ovaa, 2008). Despite it being evident that DUBs are critical to cell function, the targets and regulation processes of most DUB members have yet to be elucidated.

The largest DUB family is ubiquitin specific protease (USP), which is comprised of more than 50 USP different deubiquitinating enzymes in humans and 16 UBPs in yeast (Nijman et al., 2005). USPs have essential roles in protein regulation and function; disruption of their roles could lead to cancer progression. Thus, USP proteins have been targeted with anti-tumour chemotherapeutics (Bedford et al., 2011). As a therapeutic strategy for cancer, drugs that target DUBs are predicted to be well tolerated (Nicholson et al., 2007). The structure of multiple DUBs in the USP/UBP class including USP7, have been characterised; this has enabled identification of the method of molecular recognition (ubiquitin or ubiquitin aldehyde-complexed) used by these proteases in their active state (Saridakis et al., 2005). The ubiquitin recognition mechanism is the same for the USP/UBP class, which show homology in their catalytic sites only (Nicholson et al., 2007). and has interaction domains in their insertions and terminal extensions (Reyes-Turcu et al., 2009). These features provide USP with high selectivity for its targets.

USP7 is one member of the of USP family that due to its functional importance, has been quite extensively studied. Through its binding partners and substrates,, USP7 plays fundamental roles in diverse cellular and biological processes, such as DNA repair, cell cycle, epigenetic regulation and tumour suppression (Al-Eidan et al., 2022). Thus, mapping the USP7 network may lead to discovering new cellular functions of USP7 that are implicated in colorectal cancer. USP7 is overexpressed in many cancers, including colorectal in which it has been shown to stabilize a number of proteins that promote tumorigenesis in diverse process such as enhanced proliferation, increased angiogenesis and metastasis (Saha et al., 2023). USP7 promotes colorectal cancer (CRC) growth by upregulating various cellular pathways, including Wnt/β-catenin signalling (Li et al., 2020) and p53–Mdm2 pathway (Becker et al., 2008; Zhi et al., 2012). Finally, USP7, is one of genes identified and combined to form a prognostic mutation panel in stage II and III CRC (Sho et al., 2017).

Despite the rapidly increasing number of USP7-related studies, relatively little is known about how USP7 regulates other key cancer-related pathways such as cell-cell adhesion, an important regulator of key tumour processes including epithelial-mesenchymal transition (EMT) and metastasis. In this study, we used an inducible knockdown model of USP7 in colorectal cancer cells to identify new potential interaction partners, substrates and ultimately functions, for USP7 in colorectal cancer. To identify potential new substrates and partners of USP7, we applied a quantitative label-free LC-MS/MS proteomics approach. First, we demonstrated that the inhibition of USP7 significantly reduced cell viability, cell proliferation, and cell-cell adhesion in 2D and 3D cultures. We then selected an appropriate USP7-sensitive cell-line by mining whole genome knockout studies and used an inducible knockout of USP7 in this cell-line to perform an LC-MS/MS screen for USP7-dependent proteins. From these results, we identified the LIM domain protein Ajuba as a novel interaction partner for USP7. We then used both cell-based and molecular approaches to investigate the role of Ajuba. Using Co-IP and knockdown, we show that Ajuba interacts with USP7 and that Ajuba protein levels are dependent on USP7. With Ajuba’s known roles in cell adhesion we next showed that knockdown of either USP7 or Ajuba in dispase-treated cell-cultures significantly enhanced loss of cell adhesion. We also showed that N-cadherin and E-cadherin levels are reduced in USP7 (and partially in Ajuba) knockdown cells albeit with no evidence of direct interaction between USP7 and the cadherins. In summary, we have identified Ajuba as a novel interaction partner of USP7 that mediates USP7 functional regulation of cell-cell adhesion in colorectal cancer cells.

## METHODS

### Maintenance of cell lines

The CRC cell-line, HCT116, was cultured in McCoy’s 5A media (Life Technologies, 26600023). LS174T cells were obtained from SIGMA-ALDRICH (8706040-1VL) and were cultured in Eagle’s Minimum Essential Medium (EMEM) (ATCC® 30-2003). LS174T cells are colon carcinoma cells carrying a pTER-USP7 shRNA construct and a Tet repressor (TR)- construct (indicated as LS88) (provided by Professor Madelon Maurice, University Medical Center Utrecht) (Kessler et al., 2007). These cell lines were cultured in the RPMI media (Life Technologies, 6187010) and selected by (blasticidin 10 µg/mL (Invivo Gen, ant-bl-05) and zeocin 500 µg/mL (Invivo Gen, ant-zn-05)). Producing clones were stimulated for siRNA production by the addition of doxycycline (1 µg/mL) (Sigma, D9891-1G) for 24h, 48h and examined by Western blotting to down-regulate the endogenous USP7. DLD-1 cell lines were regularly maintained in RPMI media (Life Technologies, 6187010), whereas HEK293T Human Embryonic Kidney cells were routinely cultured in Dulbecco’s Modified Eagle’s Medium (DMEM) (Thermofisher). All cells were cultured in media supplemented with 10% fetal bovine serum and 1% streptomycin-penicillin at 37°C in CO_2_ incubator (5% CO_2_) and were regularly tested for mycoplasma. All of the colorectal cancer cell-lines used (LS88, LS174T, DLD-1, HCT116) were passaged at 70-80% confluence, washed with 1x phosphate-buffered saline (DPBS, ThermoFisher Scientific, 14190094) and detached using TrypLE™ Express (Life Technologies, 12605010) - with incubation at 37°C, 5% CO_2_ for 5 minutes. After the cells were detached, A volume of fresh media was added to the cells (1:1 ratio of media: TrypLE) and then cells were added to the new flask with an appropriate amount of fresh media.

### Protein extraction and quantification protocol

Using the Eppendorf 5415R Centrifuge at RT (room temperature), 500,000 cells were harvested and pelleted at 3,000xg for 5 minutes. Resuspension of the cell pellet in 1ml ice-cold DBS (Life Technologies, 14190094) and its centrifugation at RT thereafter was done for 5 minutes at 3,000xg. The cell pellet was re-suspended in a 1μl protease inhibitor and a 100μl RIPA buffer. After incubation at RT for 5 minutes, the supernatant collected after the lysate was centrifuged at 4°C at 16,000xg for 10 minutes. The protein in the supernatant was quantified using the Pierce BCA Protein Assay Kit (Thermo Fisher Scientific 23227) in accordance with the supplier protocol.

### RNAi knockdown

Cells were transfected according to the manufacturer (Dharmacon) protocol (Life Sciences). Cells were transfected using DharmaFect-2 (Life Sciences) with the indicated siRNA oligos (Table 1). In 6-well plates, a final concentration of 0.035μM siRNA oligo was diluted in a 1:50 HBSS (Thermo Fisher Scientific) for each well. While the DharmaFect-2 reagent was diluted in a (2:100) HBSS, the mixture was incubated for 10 minutes at room temperature. A volume of 200 μl of diluted DharamFect-2 reagent was added to each well and left for 20 minutes at room temperature. Then fresh media was made up to a total volume of 2000μl per well and then left for 72 hours in an incubator at 37°C, 5% CO_2_.

### Cellular studies with FT671 inhibitor

The USP7 inhibitor FT671 (MedChemExpress, HY-107985) was added to the cells at a final concertation of 10mM for 0h, 2h, 4h, 8h and 24h. Cells were then trypsinized and washed once with cold PBS. For cell lysate, cells were lysed in RIPA buffer (50mM Tris-HCl pH8, 150mM NaCl, 0.1% SDS, 0.5% Deoxycholic acid, 1% NP-40) supplemented with protease inhibitor (Thermo Scientific) for Western blotting.

### Immunoblotting

15μg of protein lysate was incubated in a heating block (Benchmark) at 70°□C for five minutes with sample buffer and 0.25M DTT and were loaded on 10% acrylamide gel and subjected to electrophoresis (120V, 15 minutes then 160V for 50 minutes). Using the Bio-Rad Trans-Blot Cell (Bio-Rad, 20179) proteins were transferred to a nitrocellulose membrane (GE Healthcare Life Sciences, A10224470) from the gel. (4°C, at 65V and for 2 hours) and incubated at RT for 1 hour in 10ml TBST containing 5% milk powder. Membrane was placed in its respective primary antibody solution and incubated at 4°C overnight (Table 2). Membrane segments were then washed twice with TBST and imaged using LICOR Odyssey® CLx and analysed with Image Studio Lite V5.2 software with the scan intensity set to 4. After that, images were quantified using ImageJ with the normalisation to GAPDH, actin, α-Tubulin or β-tubulin loading control to compare the density of each band.

### Cell viability

Attempts have been made to evaluate the effect of 0.1 ul doxycycline treatment on LS88 cells during the proposed time course through CellTitre assay (Promega, 0000333156). Cells are treated using control growth media or (100 ul) growth media supplemented with 0.1ul doxycycline time points of 24 hours, 48 hours, and 72 hours, respectively. The measurement of cell viability took place using Promega Glomax Multi detection system. Similarly, the number of cells was determined with standard column calculations and change was calculated in comparison to the cell’s growth in control growth media.

### Colony formation

USP7 knockdown was performed using small interfering ribonucleic acid (siRNA) oligo; 2000 cells were plated in 6 well plates and the cells were incubated for 7-14 days. The cells were washed with PBS after the incubation and fixed with 4% paraformaldehyde (PFA) (4g PFA + 80 ml + 20 ml of 1X PBS) for 30 minutes. After the fixing, the cells were washed gently with dH2O twice and stained with 0.5% crystal violet and 20% methanol for 30 minutes. The cells were washed after staining with dH_2_O three times and left to dry in the hood overnight before the imaging and counting took place. Colonies with at least 50 cells were counted. Each experiment was performed in triplicate.

### Spheroid assays and quantifications

After siRNA transfections were performed in two-dimensional (2D) cultures and left 96 hours post-transfections, cells were cultured in a 96-well ultralow attachment plate (Corning-3474) in 100µl with plating densities of 1000 (HEK 293T), 3000 (LS88 and LS174T) and 4000 (DLD-1) cells/well. Cells were cultured in 1:1 DMEM: F12, (Gibco® by Life Technology) media plus 10% FBS, and 1% Pen Strep (Gibco® by Life Technology). The cells were incubated at 37°C and 5% CO_2_ for 14 days. Then, the images were taken using ×40 magnification. The volumes of the spheroids were calculated using the formula of volume = (4/3)πr3. ImageJ (version1.42q) was used to determine the volume of a spheroid. CellTiter-Glo® cell viability assay was performed with the addition of 100 µl of CellTiter-Glo® reagent into each well and incubated at room temperature for 1h, followed by measurement using GloMax® Discover Microplate Reader (Promega).

### Immunofluorescence microscopy

Sterile glass coverslips (22X22) were used to grow HCT116 knockout USP7 Cells into cell culture 6-well plate and the cell fixed using 4% Paraformaldehyde and quenched using quench formaldehyde with 10 mM Glycine treatment. Cells were then permeabilised with 0.2% Triton100 and blocked 5 min at RT in 3% BSA Blocking solution (PBS+ Tween 20). Next, the slides were incubated with primary antibodies diluted in 5% BSA solution for a time period of 1 hour at RT. Slides were then incubated with secondary antibodies (Alexa fluor) for 1 hour 1:1000. The slides were then rinsed with BSA solution then with PBS. Coverslips were mounted overnight with the help of ProLong Gold antifade mounted with DAPI. A Leica TCS SP8 microscope was then made use of to visualize the slides; this was acquired using LAS X Core software (V3.3.0). For the sub-cellular analysis of LS88, cells were untreated / treated with doxycycline. Fiji (ImageJ) was used to create Regions of Interest for the measurement of image mean grey value for cytoplasm, nucleus and local background (n=20). Data were corrected by subtracting local background values from cytoplasm and nucleus mean grey values. Mean, Standard deviation and T-Test were calculated, and intensity data was expressed as percentage of control values.

### Immunoprecipitation

The cells were lysed for 30min at 4 °C in pNAS buffer (50 mM Tris/HCl (pH 7.5), 120 mM NaCl, 1 mM EDTA and 0.1% Nonidet P-40) with protease inhibitors. Cells were scratched and kept on ice for 30 minutes, and then proteins were quantified using bicinchoninic (BCA) assay. For input, 40 µg of samples were aliquoted. Protein G Sepharose beads were utilised after washing them with 1X phosphate-buffered saline (PBS) three times. The concentration of immunoglobulin G (IgG) agarose and the antibody samples were 1 mg/ml. In addition, 40 µl of the shacked beads were added to the IgG and antibody samples and rotated for one hour. Next, the IgG agarose and antibody samples were spun down in the centrifuge (570 x g at 4°C) for 1 minute. In a new Eppendorf, 950µl from the supernatant was taken. Subsequently, 50 µl of the Protein G Sepharose beads were added to the antibody samples and IgG. Indicated antibodies and IgG were added to the lysate for 16 h at 4 °C. The samples and IgG were spun down (570 x g at 4°C for 1 minute). Immunoprecipitates were washed four times with cold PBS followed by the addition of sodium dodecyl sulphate (SDS) sample buffer. The bound proteins were separated on SDS polyacrylamide gels and subjected to immunoblotting with the indicated antibodies.

### RNA extraction

RNeasy® Mini Kit (Qiagen, UK) was used to extract RNA from the samples according to the manufacturer’s protocol. Cells plated in a 6-well plate were washed with 1X PBS twice after the media was removed. The cells were lysed with 350 µl of Buffer RLT in each well. After pipetting several times, 350µl of 70% ethanol were added into the lysate and mixed well. 700 µl of mixture was then transferred into each RNeasy Mini spin column and centrifuged at maximum speed for 30 seconds. After discarding the flow-through, 500 μl of Buffer RW1 was added onto the spin column and centrifuged, following two steps using 500 μl of the Buffer RPE and centrifuged at maximum speed for 30 seconds and 2 minutes. Finally, 40 µl of RNase-free water was added to RNeasy spin column which was placed in a new 1.5 ml Eppendorf and centrifuged at maximum speed for 1 minute. To quantify RNA concentration, a Nanodrop Spectrophotometer 2000c, (Thermo Fisher Scientific, UK) was conducted. And the RNA samples were stored at -80°C until qPCR was performed.

### Quantitative real-time PCR

RNA samples with a final concentration of 20 ng/μl were diluted in RNase free water (Qiagen, Germany) and used in this experiment. A QuantiNova SYBR Green RT-PCR kit (Qiagen, Germany) was utilised to perform real-time PCR according to the manufacturer’s instructions. The reaction set-up (Table 3), the primers (FW-USP7 - 5’-GGA CAT GGA TGA CAC CA -3’), (RV-USP7- 5’-TCA CTC AGT CTG AAG CG -3’), (FW-AJUBA- 5’-GAG AAC CCT CGG GGA TTG A -3’) and (RV-AJUBA- 5’-GCA CTT GAT ACA GGT GCC GAA -3’) (Integrated DNA Technologies), and the thermocycling conditions were listed (Table 4). Work on each sample was performed in triplicate in a 96-well plate on a StepOnePlusTM Real-Time PCR System (ThermoFisher Scientific, United Kingdom). The fluorescent dye was released during amplification. The point at which the intensity exceeded the threshold was recorded in order to determine the Ct value. The more significant the Ct value, the later the intensity exceeded the threshold, and the lower the expression of that specific gene. Gene expression of each sample was evaluated from the ΔΔCt value utilising ACTB (β-Actin) as a control. The results were analysed in GraphPad Prism, version 9.3.1 (GraphPad Software Inc, CA, United States).

### Dispase mechanical stress assay

Intercellular adhesion was evaluated by incubation with the dispase enzyme on cell monolayers followed by a mechanical stress test, then the cell monolayers were assessed using light microscopy. LS88, control siRNA, and USP7- or Ajuba-depleted cells were seeded in 6-cell culture plates (Corning) with three replicates for each condition and grown until reaching confluency. The cell monolayers were washed with Hanks’ Balanced Salt Solution (HBSS) (14025092, ThermoFisher Scientific) after the media was removed from the cultures. The cell monolayers were incubated with dispase enzyme (2.5 units/ml in HBSS) (D4693, Sigma) at 37 °C for 30 minutes or longer in order to detach the monolayers from the plate bottom. Once the monolayers detached, they were imaged using a Leica M2 16 F microscope and S Viewer V3.0.0 software. After that, the monolayers were subjected to mechanical stress by pipetting with 1ml HBSS and shaking in the shaker for 3 minutes at 300 rpm. The fragments were counted for all three conditions and imaged.

### Experimental Design and Statistical Rationale

This study compares the proteomes of LS88 cells with USP7 inducibly knocked-down using doxycycline to control cells with wild-type levels of USP7. We analysed 9 mass-spectrometry samples in total consisting of 3 groups with 3 biological replicates in each group (1) LS88 untreated at 0 hours, (2) LS88 untreated at 72 hours and (3) LS88 treated with doxycycline at 72 hours. These treatments were based upon results presented in Figure 1 and in the Supplementary Data, and we observed good correlation between biological replicates based upon the PCA and unsupervised clustering shown in Figure 2. To identify proteins that are significantly decreased following treatment with doxycycline and USP7 knockdown, we compared the LS88 cells untreated (at 72hrs) with LS88 cells treated with doxycycline (at 72hrs). Only proteins identified with peptides across all 6 of these samples were considered when performing the comparisons. For all analysis of Western blot densitometry, paired or unpaired Student’s t tests were performed as appropriate in GraphPad Prism 9.2 software. For all statistical comparisons of more than 2 groups, ordinary one-way analysis of variance (ANOVA) was conducted with GraphPad Prism 9.2 software.

**Figure 1.**
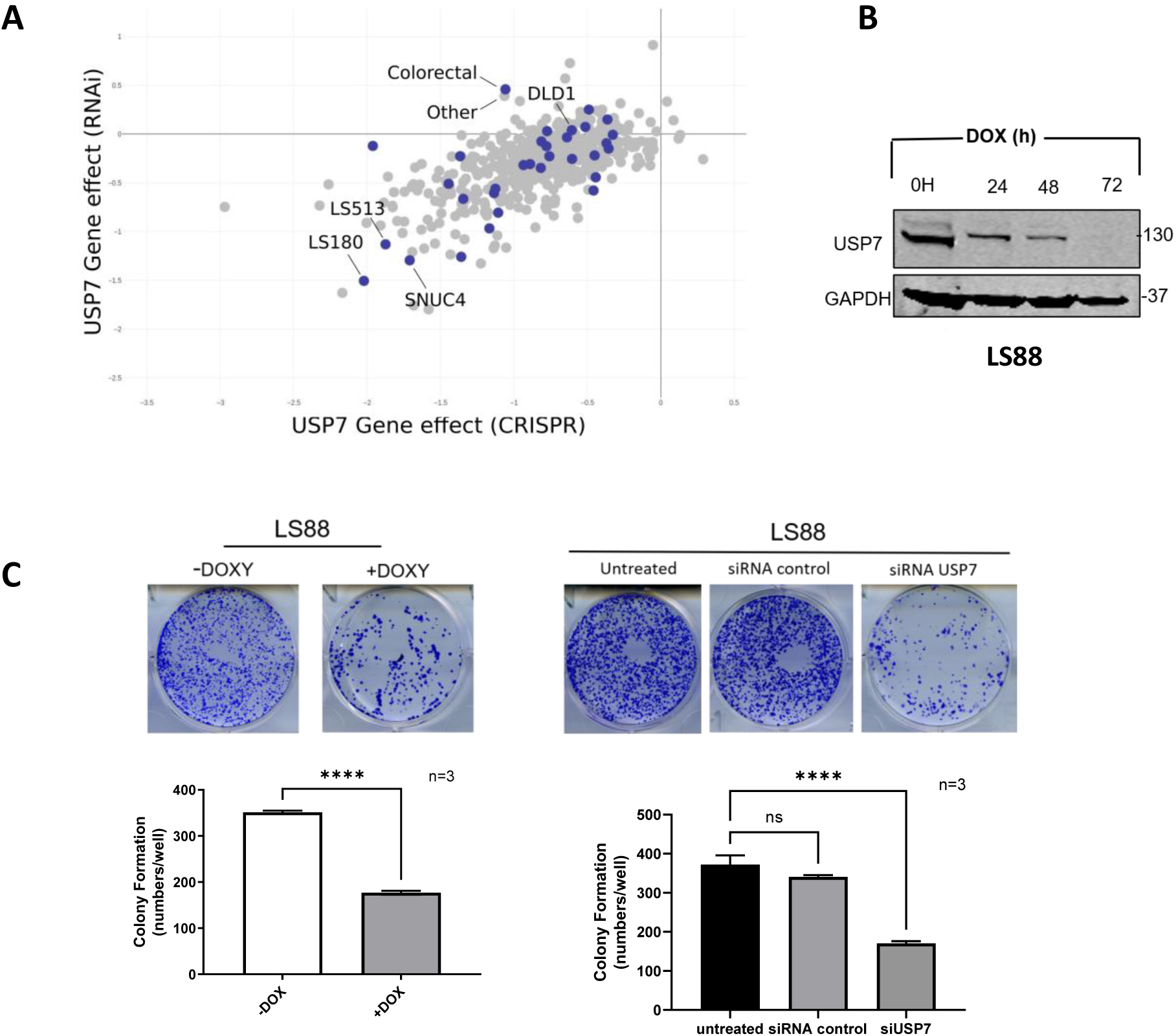
LS180 is an effective model for investigating USP7 function in colorectal cancer. **(A)** Scatterplot comparing the distribution of USP7 gene effect processed scores (CERES, DEMETER2) between the RNAi dataset (y) and CRISPR dataset (x). Selected cell-lines with the largest USP7 gene-effect scores from RNAi and CRISPR are labelled in addition to DLD1, an additional CRC cell-line used in this study. All analysis performed in DepMap portal (depmap.org) **(B)** Efficient degradation of USP7 at the indicated time points. LS88 colon carcinoma cells stably carrying a pTER-USP7 shRNA construct, and a Tet repressor (TR)-construct were treated with 1 µg/mL doxycycline for 24, 48, and 72 hours and examined by Western blotting for reduction of endogenous USP7, GAPDH was used as a loading control. (C) LS88 cells (with inducible siRNA targeting USP7) were treated with 1 µg/mL concentrations of doxycycline for 72h before being seeded in 6 well plates and incubated for 7-14 days. Graph represents mean ± SD (two tailed, unpaired t-test), P ≤ 0.0001., LS88.

**Figure 2.**
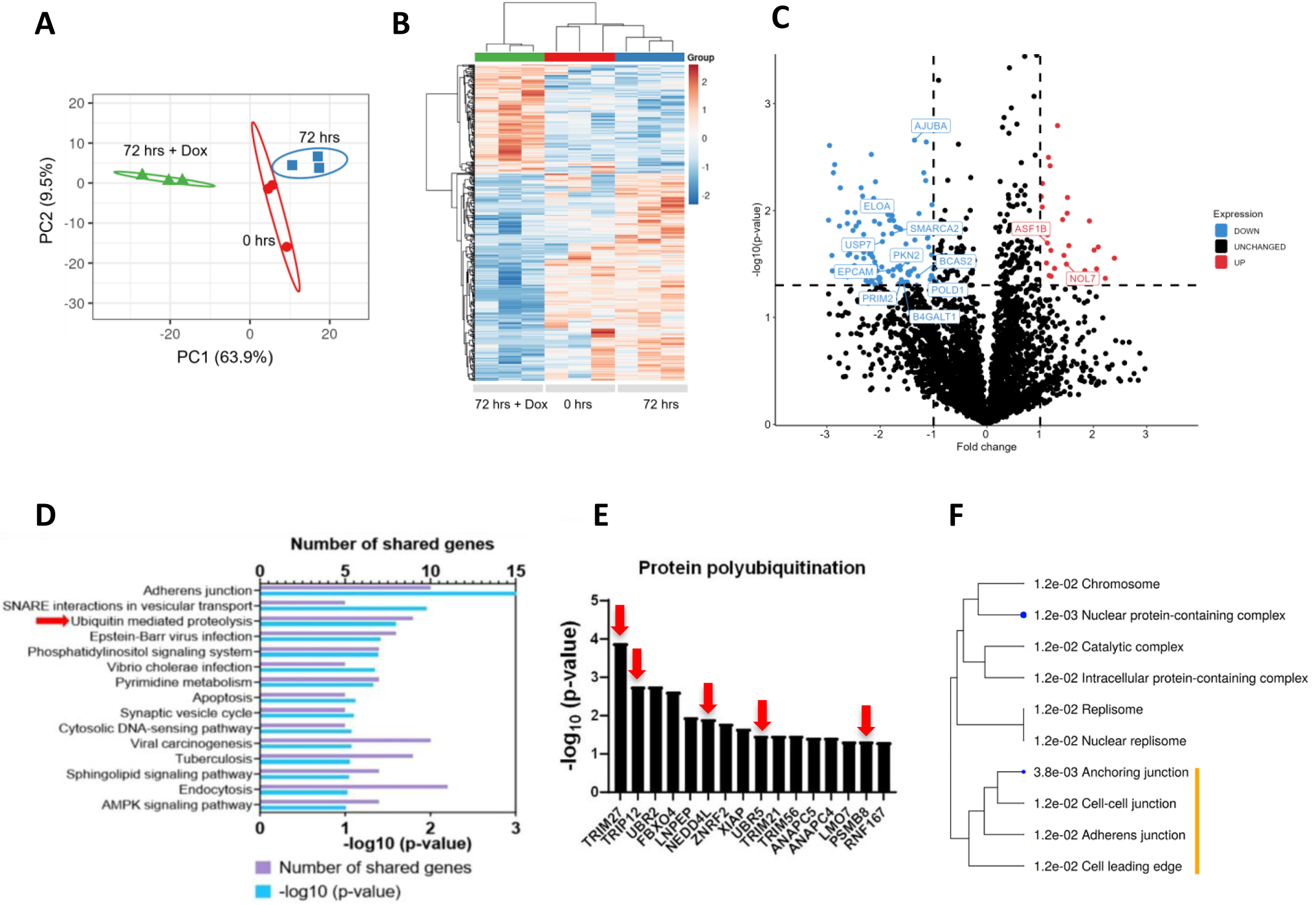
Proteomic analysis of LS88 with/without doxycycline treatment for USP7 knockdown. (A) PCA scores plot showing the clustering of proteins according to their distributions in different groups. Control LS88 represents cells at hour zero (0 hrs), LS88 treated with doxycycline for 72 h (72 hrs + doxycycline) or left without treatment for 72 h (72 hrs). (B) Heatmap of normalised intensity values (light/low – dark/high) for each group (0 hrs untreated; 72 hrs untreated; 72 hrs + doxycycline). All proteins (N = 444) with p < 0.05 for treated vs. untreated (at 72 hrs) are included. (C) Volcano plot showing differentially responsive proteins in comparison of 72hrs untreated vs 72hrs doxycycline-treated cells. Selected proteins from enriched pathway and process categories are labelled. (D) Annotation analysis of the sets of proteins (p<0.05) decreased following doxycycline treatment (after 72hrs). Pathways from DAVID tool are sorted by p value and the top 15 most significant pathways shown. Ubiquitin-mediated proteolysis category is indicated by arrow. (E) The top 15 most significantly decreased proteins (p< 0.05) following doxycycline treatment (after 72 hrs) that are involved in protein polyubiquitination. TRIM27, a USP7 protein-complex partner, is the most significantly altered protein following USP7 knockdown in this category (Analysis performed in Metascape). Proteins that are known to be direct substrates or interaction partners of USP7 (according to STRING) are indicated with arrow. (F) Dendrogram showing significant Gene Ontology cellular components for the set of proteins (p<0.05) decreased following doxycycline treatment. Enrichment p-values are indicated and only GO terms with FDR p-value < 0.05 were included. GO terms associated with cell-cell junction functions are highlighted.

### Proteomic sample preparation and mass-spectrometry

Lysis of the protein pellets (0.1M TEAB, 0.1% SDS) was performed with pulsed sonication, and then samples were centrifuged (up to 10 minutes, 13,000 x g). The extraction of methanol/chloroform was done on 40μg of protein for each lysate and finally, the pellets were re-dissolved in 100μl of 6M Urea (Sigma), 2M thiourea (Sigma), 10mM HEPES buffer (Sigma), ph7.5. This was followed by the reduction of the samples (1M DTT), alkylation (5.5M iodoacetamide), and then dilution in 400µl of 20mM ammonium bicarbonate (Sigma) and later digested alongside trypsin (Promega) (1/50 w/w) overnight. Acidification of each sample <3.0 was done with the trifluoroacetic acid (TFA) (Sigma) and solid phase extraction performed using waters oasis HLB prime μElution 96-well solid phase extraction plate. Plate collection vessel was used in the plate vacuum manifold and the plate was conditioned with the addition of 200ul of acetonitrile. The wells were equilibrated through 200μl of 0.1 % TFA and then inserting a 96 well collection plate. There was a further application of 100μl of sample to each well and of a low vacuum. A protective cover was used to cover the 96 well collection plate, and was stored at 4 ℒC. The collection plate was replaced with the vessel again. The wells were washed using 2X 700μl of 0.1 TFA and then 200μl of water in order to remove buffer and salts. Addition of 70% of acetonitrile to the well was done to evaporate the remaining liquid followed by re-suspension in buffer A (H_2_O + 0.1% Formic Acid). To meet the internal quantification standard, 200 fmol of Waters enolase (Saccharomyces cerevisiae) standard were further added to each of the samples.

Peptide extracts (1 μg on column) were separated on an Ultimate 3000 RSLC nano system (Thermo Scientific) using a PepMap C18 EASY-Spray LC column, 2 μm particle size, 75 μm x 75 cm column (Thermo Scientific) over a 140 min (single run) linear gradient of 3–25% buffer B (0.1% formic acid in acetonitrile (v/v)) in buffer A (0.1% formic acid in water (v/v)) at a flow rate of 300 nL/min. Peptides were introduced using an EASY-Spray source at 2000 V to a Fusion Tribrid Orbitrap mass spectrometer (Thermo Scientific). The ion transfer tube temperature was set to 275 °C. Full scans were acquired in the Orbitrap analyser using the Top Speed data dependent mode, performing a MS scan every 3 second cycle, followed by higher energy collision-induced dissociation (HCD) MS/MS scans. MS spectra were acquired at resolution of 120,000 at 300 m/z, RF lens 60% and an automatic gain control (AGC) ion target value of 4.0e5 for a maximum of 100 ms. Peptide ions were isolated using an isolation width of 1.6 amu and trapped at a maximal injection time of 120 ms with an AGC target of 300,000. Higher-energy collisional dissociation (HCD) fragmentation was induced at an energy setting of 28 for peptides with a charge state of 2–4. Fragments were analysed in the orbitrap at 30,000 resolution.

### Mass-spectrometry data processing and analyses

Analysis of raw data was performed using (1) Proteome Discoverer software (Thermo Scientific) (version 2.4) and (2) Mascot search engine (version 2.7.0.1). For (1) the data was processed to generate reduced charge state and deisotoped precursor and associated product ion peak lists. These peak lists were searched against the Human UniProt protein database (080620) (570,420 protein sequences) A maximum of one missed cleavage was allowed for tryptic digestion and the variable modification was set to contain oxidation of methionine and N-terminal protein acetylation. Carboxyamidomethylation of cysteine was set as a fixed modification. The false discovery rate (FDR) was estimated with randomized decoy database searches and were filtered to 1% FDR.

For Mascot searches, peak lists were generated using ProteoWizard’s msconvert (version 3.0.23213) to convert the raw peak .RAW files into .mgf peak lists for Mascot (2023 release). Thereafter, the .mgf files searched using Mascot server using its default parameters against UniProt’s *Homo sapiens* proteome (UniProt proteome ID UP000005640) and Proteomics Identifications Database (PRIDE) contaminants spectra (Release: 2015-04-01, Version: 2015-04). The false discovery rate threshold was set at 0.05. Trypsin (X-[K/R]) was used for the digestion with two allowable missed cleavages. The variable modifications used in the search included: oxidation of methionine, and carbamidomethylating of cysteine. No fixed modifications were included in the search. Mass tolerance for precursor ions was set at ± 1.2 Da and for fragment ions at ± 0.6 Da. Identified proteins were filtered by setting a false discovery rate threshold of 1%.

The mass-spectrometry data has been submitted to the PRIDE database with the following details:

Project accession: PXD039488

Project DOI: 10.6019/PXD039488

Reviewer account details:

Username

Password: ReyeLEaT

### Additional bioinformatics analyses

For the analysis of vulnerabilities across cancer cell lines, the Cancer Dependency Map (Tsherniak et al., 2017) CRISPR and RNAi data were used (version Public 24Q2). All analysis and plots were generated through the DepMap portal (https://depmap.org/portal). To investigate pathways and gene enrichment in the mass spectrometry (MS) proteomics data, the set of 445 proteins which were significantly (p < 0.05) altered between the doxycycline treated LS88 cells and the untreated LS88 cells at 72hrs, were analysed. Several tools were used: the (DAVID) functional annotation web tool (version 6.8; https://david.ncifcrf.gov) (Huang et al., 2008), the ShinyGO interface and tool (Ge et al., 2020) (default settings, FDR cutoff = 0.05), and Metascape pathways analysis tool (Zhou et al., 2019) (default settings).

## RESULTS

### Selection and optimization of a cell model for analysis of USP7 function in colorectal cancer

To identify a suitable cell model for analysis of USP7 function in colorectal cancer, we first analysed publicly available cancer-dependency profiles from the DepMap project (Tsherniak et al., 2017). The DepMap project includes both large-scale CRISPR and RNAi screens of many cancer-cell lines and provides a significant resource for identifying the vulnerabilities of different cancer types using knockout or knockdown screens. We analysed the CRISPR and RNAi dependency scores across all cell-lines in the DepMap as shown in Figure 1A and noted that several colorectal cancer cell-lines show strong negative gene-effect scores in both the CRISPR and RNAi datasets. Notably, growth of the LS180 cell-line, is significantly inhibited through either USP7 knockdown or knockout in the DepMap project (Figure 1A). LS180 is a human colon adenocarcinoma cell line derived from a 58-year-old Caucasian female with Dukes type B adenocarcinoma of the colon (ECACC 87021202). The related LS174T cell line (ECACC 87060401) was derived from the same patient (“ECACC General Cell Collection: 87021202 LS180,” n.d.), and were subsequently modified to express a doxycycline-inducible siRNA construct pTER-USP7 (cell-line known as LS88) (Kessler et al., 2007). We used this LS88 cell-line as the basis of our proteomic screen in this study. To identify suitable conditions for the proteomic screen, we confirmed that treatment of LS88 cells with doxycycline induced knockdown of USP7 in a time-dependent manner (Figure 1B) and that levels of Trim27 and p53, two key proteins regulated by USP7 were regulated as expected according to previous studies (Meulmeester et al., 2005) following doxycycline treatment (Supplementary Figure S2). We next showed that cell viability is significantly affected following treatment of LS88 cells with doxycycline or siRNA induced knockdown of USP7 (Figure 1C). Further results of cell viability and colony formation following USP7 knockdown are provided in Supplementary Figure S1.. We also confirmed that the observed decreases in cell viability were due to USP7 knockdown and not treatment with doxycycline per se, since doxycycline is known to have a negative effect on invasive potential and cell proliferation across varied human CRC cell lines (Onoda et al., 2004; Sagar et al., 2010) (Supplementary Figure S3). Finally, we also showed that inducible knockdown for 72 hrs in LS88 cells with doxycycline has similar effects to USP7 depletion via siRNA in another colorectal cancer cell-line, DLD-1. Notably, the DLD-1 cell-line, in common with many other colorectal cancer cell-lines has mutant truncated APC, and a previous study indicated that USP7 has a specific role in maintaining Wnt signalling in tumours with APC mutations (Novellasdemunt et al., 2017). We also observed that in the DepMap (depmap.org) gene-effect data, the effect of USP7 knockout or knockdown is markedly less pronounced in colorectal cancer cell-lines with APC mutation than in those without (Supplementary Figure S4). In summary, the USP7 inducible knockdown cell-line LS88 in the highly USP7-dependent CRC cell line of LS180 represent an appropriate model for studying USP7 function in colorectal cancer. The effects are comparable to other widely used CRC cell-lines following USP7 depletion.

### Differential analysis of the USP7 network using inducible knockdown proteomics

To broadly characterize proteome-wide altered protein expression in response to USP7 knockdown, we performed whole-cell profiling using quantitative label-free LC-MS/MS. Based on the altered phenotypic properties of the doxycycline-treated LS88 cells observed above, we compared doxycycline-treated LS88 cells at 72hrs to control (untreated) cells at 0 and 72 hrs after the start of the experiment. Three replicate samples were profiled using LC-MS/MS for each group. Data-dependent quantitative mass spectrometry analysis identified a total of 7,301 proteins of which 445 proteins were found to be significantly (p < 0.05) differential between the treated and untreated samples at 72hrs. We then analysed the sample reproducibility as shown in the PCA and heatmap analyses and volcano plot in Figure 2A-C. Since our goal is to identify proteins whose expression decreases alongside depletion of USP7 (and are therefore potential substrates of the USP7 deubiquitinase), we analysed the set of 293 proteins which were significantly (p < 0.05) decreased following doxycycline treatment. We first analysed these proteins using DAVID (Huang et al., 2008) to broadly assess the most significant pathways (Figure 2D). We noted that proteins related to ubiquitination-mediated proteolysis represent a significantly enriched group and show this set of proteins in Figure 2E. Notably, the top 2 proteins, TRIM27 and TRIP12 are known interaction partners of USP7 ((Liu et al., 2016; Zaman et al., 2013), suggesting that our dataset could be a rich source of USP7 substrates and novel interaction partners. We also noted that from the DAVID analysis, adherens junctions proteins represent the most significantly enriched group within the data. We further analysed this finding using ShinyGo (Ge et al., 2020) to represent the 10 most significant GO cellular components as shown in Figure 2F. Although USP7 has been implicated in the regulation of several cancer-associated pathways, its role in the regulation of cell-cell adhesion is relatively unstudied. We focused on the Ajuba protein, that was previously found to promote growth, migration and metastasis in colorectal cancer, in addition to its roles and functions in regulating cell-cell adhesion and pathways such as Hippo (Jia et al., 2020). Finally, we annotated the volcano plot (Figure 2C) with key proteins from this study and with selected proteins from the most significant GO cellular component categories in Figure 2F. Finally, protein intensity values are provided for those proteins analysed by Western blot in Supplementary Figure S5.

### USP7 sustains Ajuba protein stability in CRC cells

Ajuba was significantly decreased in response to doxycycline in the MS proteomic analysis, and we first confirmed this response using immunoblotting. The level of Ajuba protein after USP7 inhibition was determined at different time points (0, 24, and 72hr) using doxycycline (1 µg/mL) in LS88. We observed a reduction of Ajuba expression after USP7 depletion, and a particularly marked decrease in Ajuba after 72hrs following doxycycline treatment, comparable to the time point used in the MS proteomics study (Figure 3A, 3B). We further explored this effect by using FT671, a non-covalent selective USP7 inhibitor. We treated LS88 cells with the FT671-USP7 and the levels of Ajuba was analysed using Western blot (Figure 3C). As the figure demonstrates, the Ajuba protein levels were changed after 48, 72, and 96 hours after the treatment. Similarly, the results from a different colorectal cancer cell line (HCT116) showed a reduction of Ajuba protein expression after the cells were treated with the inhibitor for 48 and 72 hours (Figure 3D).

**Figure 3.**
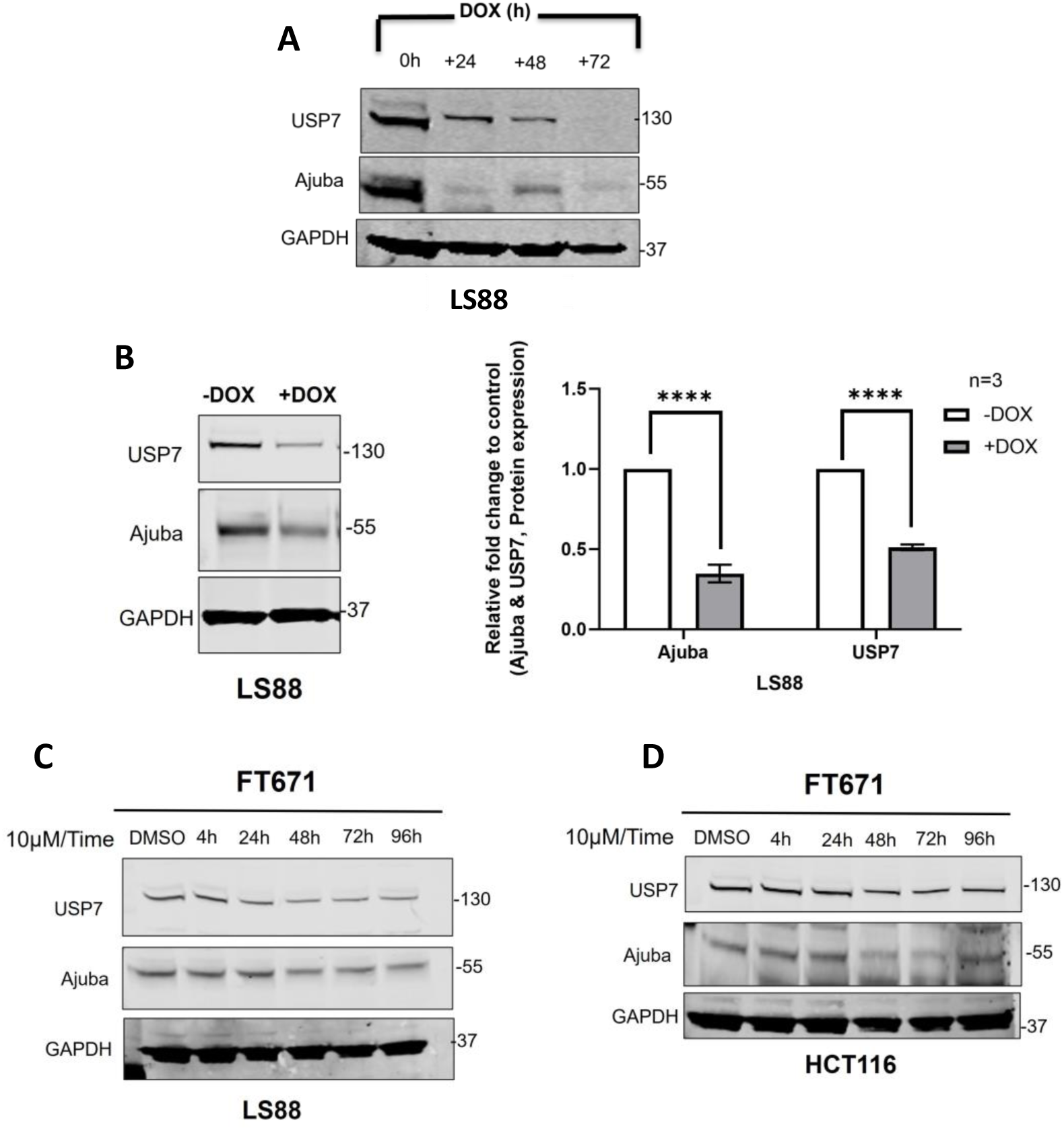
Western blot analysis of Ajuba abundance following USP7 knockdown. **(A)** Total cell lysate treated with doxycycline in different time point were collected and analysis using immunoblotting**. (B**) LS88 cells treated or untreated with doxycycline were examined on Ajuba protein expression. Unpaired Student’s t test was used for three independent experiments. P < 0.0001. GAPDH was used as a loading control. **(C)** Total cell lysates from LS88 and **(D)** HCT116 treated with FT671 (10 μ M) for indicated times. GAPDH is a loading control.

Next, we analysed the sub-cellular localisation of Ajuba and USP7 using immunofluorescence.. As shown in (Figure 4-B) Ajuba and USP7 protein levels were markedly decreased in LS88 cells treated with doxycycline (1 µg/mL) where USP7 was downregulated. Ajuba was previously reported to localize mainly in nucleoplasm & the Golgi apparatus (Thul et al., 2017), and USP7 was reported to localize predominantly to the nucleus (Zapata et al., 2001). We observed that Ajuba and USP7 protein levels were very significantly (p< 0.0001) higher in nuclei than in cytoplasm (p< 0.001) in the cells treated with doxycycline which is consistent with their localization (Figure 4C).

**Figure 4.**
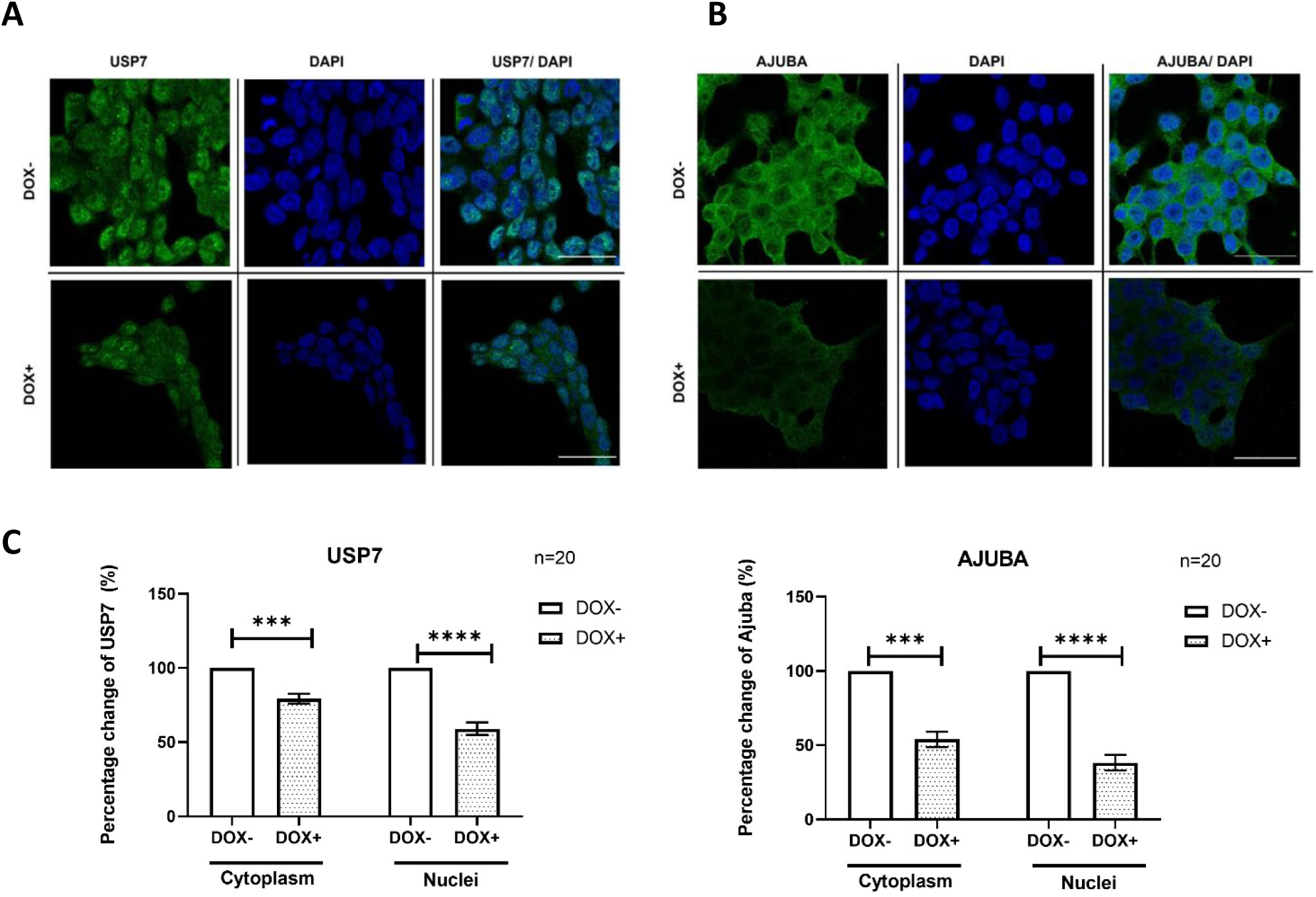
Immunofluorescence analysis of Ajuba levels following USP7 knockdown in LS88 cells. **(A),** Immunofluorescence staining of Ajuba **(B),** USP7 (green) in LS88 cells treated with doxycycline (DOX+) or untreated (DOX−). Cells were fixed and stained with DAPI (blue) to stain nuclei. Scale bars: 30 μm. **(C),** Data were corrected by subtracting local background values from cytoplasm and nucleus mean grey values. Statistical differences were determined by Unpaired t-test of percentage change of Ajuba and USP7 in cytoplasm and nuclei in LS88 cells. Mean ± SD.

In order to further investigate the regulatory relationship between Ajuba and USP7, we used qRT-PCR to show that Ajuba mRNA expression was unaffected by USP7 downregulation in doxycycline treated LS88 cells (Figure 5A-B), showing that USP7 does not regulate Ajuba at the transcriptional level. We next investigated whether Ajuba is a protein-protein interaction partner of USP7 by using immunoprecipitation with anti-USP7 antibodies in LS88 cells. We found that USP7 interacts with Ajuba in colorectal cancer cells when USP7 is pulled down and analysed by immunoblotting (Figure 5C), showing that Ajuba is indeed a physically associated interaction partner of USP7 in colorectal cancer. We also found in both LS88 and LS147T cells that reciprocal knockdown of Ajuba does not alter USP7 protein levels (Figure 5D), indicating that USP7 and Ajuba are not co-complexed proteins with mutual regulation of protein stability, but rather that Ajuba stability is dependent on USP7 but not vice-versa.

**Figure 5.**
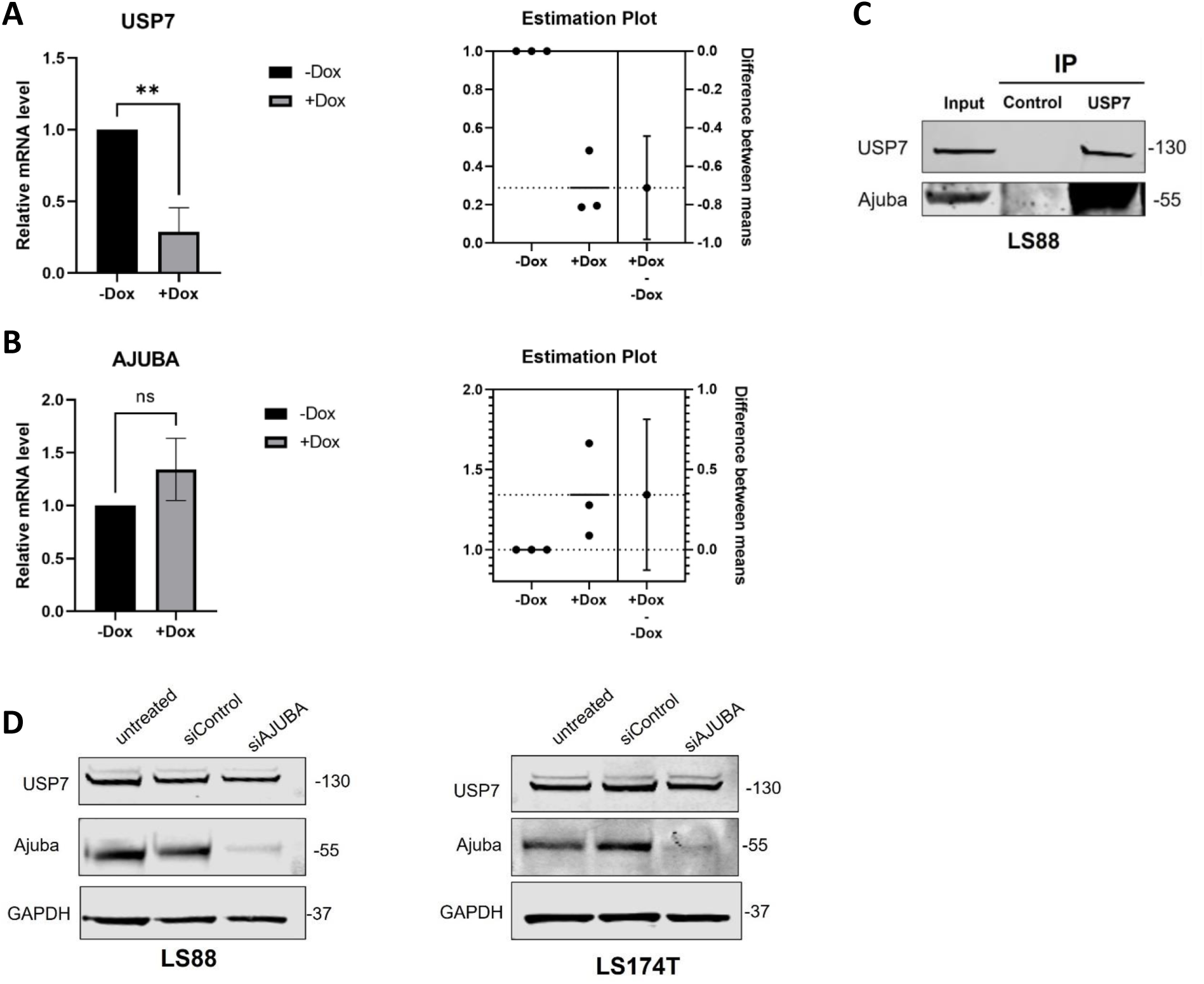
USP7 interacts with Ajuba and Ajuba knockdown does not affect USP7 protein levels in CRC cells. **(A)** mRNA levels of USP7 in doxycycline-treated (DOX+) or untreated (DOX-) LS88 cells. **P < 0.01. Data are mean ± SD. **(B)** mRNA levels of Ajuba in DOX+ or DOX-LS88 cells. Unpaired t-tests were used for analysis in order to assess any differences between the means. Experiments were performed in triplicate, n=3. **(C)** LS88 cells total lysates were immunoprecipitated with anti USP7 antibody or IgG, and Ajuba was analysed using indicated antibodies in LS88 cells. **(D)** LS88 and LS174T cells expressing Ajuba, control siRNA or Ajuba siRNA were analysed by western blotting and USP7 protein expression were examined using the indicated antibodies. As a loading control, GAPDH protein levels were assessed.

### Knockdown of USP7 or Ajuba causes disruption of cell-cell adhesion in CRC cells

Since cell-cell and adherens junctions were found to be one of the most significantly enriched functions from the proteomic analysis (Figure 2), we analysed whether knockdown of USP7 or Ajuba resulted in phenotypic changes to cell-cell adhesion. Dispase assays allow for the separation of cell culture monolayers that can then be subjected to mechanical stress. As shown in Figure 6, mechanical stress of siUSP7 or siAJUBA treated cells led to significantly increased fragmentation of monolayers as compared to control siRNA treatments, showing that reduction of either USP7 or Ajuba causes a very significant reduction in the mechanical strength of cell-cell adhesion in CRC cells.

**Figure 6.**
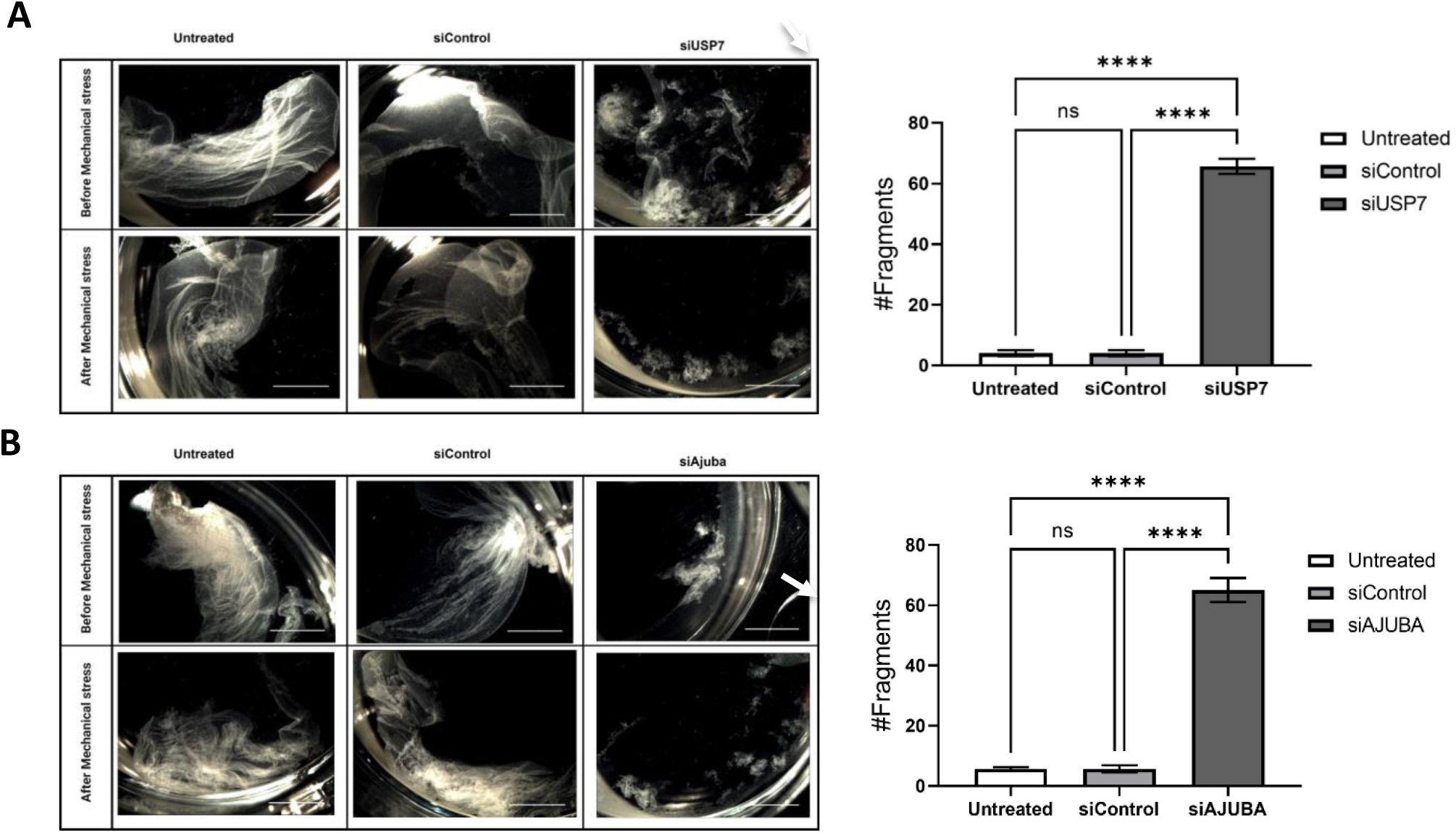
Mechanical stress analysis of cell-cell adhesion in LS88 cells. **(A)** Monolayer cultures untreated or treated with siControl or siUSP7 were treated with dispase and incubated until the monolayer was detached from the culture surface, followed by shaking to induce fragmentation. The fragments were counted for all three conditions and the bar graph shows the average number of fragments. One-way ANOVA was used. **** P < 0.0001. Mean ± SD from three independent experiments. Arrow indicates increased fragmentation of monolayers following mechanical stress. Scale bar 5mm. **(B)** Monolayer cultures untreated or treated with siControl or siAJuba were treated with dispase and incubated until the monolayer was detached from the culture surface, followed by shaking to induce fragmentation. The fragments were counted for all three conditions and the bar graph shows the average number of fragments. One-way ANOVA was used. **** P < 0.0001. Mean ± SD from three independent experiments. Arrow indicates increased fragmentation of monolayers following mechanical stress. Scale bar 5mm.

### Knockdown of USP7 or Ajuba disrupts cadherin but not catenin protein levels

To identify the underlying mechanisms of how cell-cell adhesion is disrupted through USP7 or Ajuba knockdown, we examined the expression of cell junction associated proteins. Since Ajuba has been identified to interact with several cadherins and catenins (CDH1/E-cadherin, CTNNA1/ α-catenin, CTNNB1/ β-catenin, CTNND1/ δ-catenin and JUP/plakoglobin), we focused our analysis on this group of proteins. Using immunoblotting, we investigated the protein levels of multiple cadherins and catenins following USP7 or Ajuba knockdown. As shown in Figure 7A, both N-cadherin and E-cadherin protein levels were reduced after USP7 inhibition in LS88 cells, and similarly the expression of N-cadherin and E-cadherin were reduced following USP7 knockdown in HCT116 cells (Figure 7B). We also observed that treatment of LS88 and HCT116 cells with the USP7-inhibitor FT671 across a time course resulted in decreased N-cadherin expression (Figure S6). Since these findings suggest that USP7 may regulate cadherin expression in CRC cells, we employed IP-coupled westerns (co-IP) to determine whether there is a physical interaction between USP7 and E-cadherin. This analysis (Figure 7D) did not indicate that there was a physical interaction between the proteins, suggesting that the reduction in E-cadherin expression is not mediated via interaction with USP7 and that E-cadherin is not a substrate of the deubiquitinase. Although USP7 has been shown to regulate Wnt/ β-catenin signaling in CRC (Herbst et al., 2014; Ji et al., 2019; Novellasdemunt et al., 2017), we found no altered β-catenin protein expression following USP7 knockdown in multiple CRC cell-lines (Figure S7), except in SW480 cells, that carry the APC truncation mutation which results in β-catenin stabilisation, in line with a previous publication (Herbst et al., 2014). Similarly, there was no effect on either α-catenin or JUP/plakoglobin following USP7 knockdown (Figure 7C). We observed that levels of JUP were increased (p< 0.05) following USP7 depletion in the proteomics data, but did not observe any increase using Western blots, nor was there an interaction detected via co-IP (Figure 7D)).

**Figure 7.**
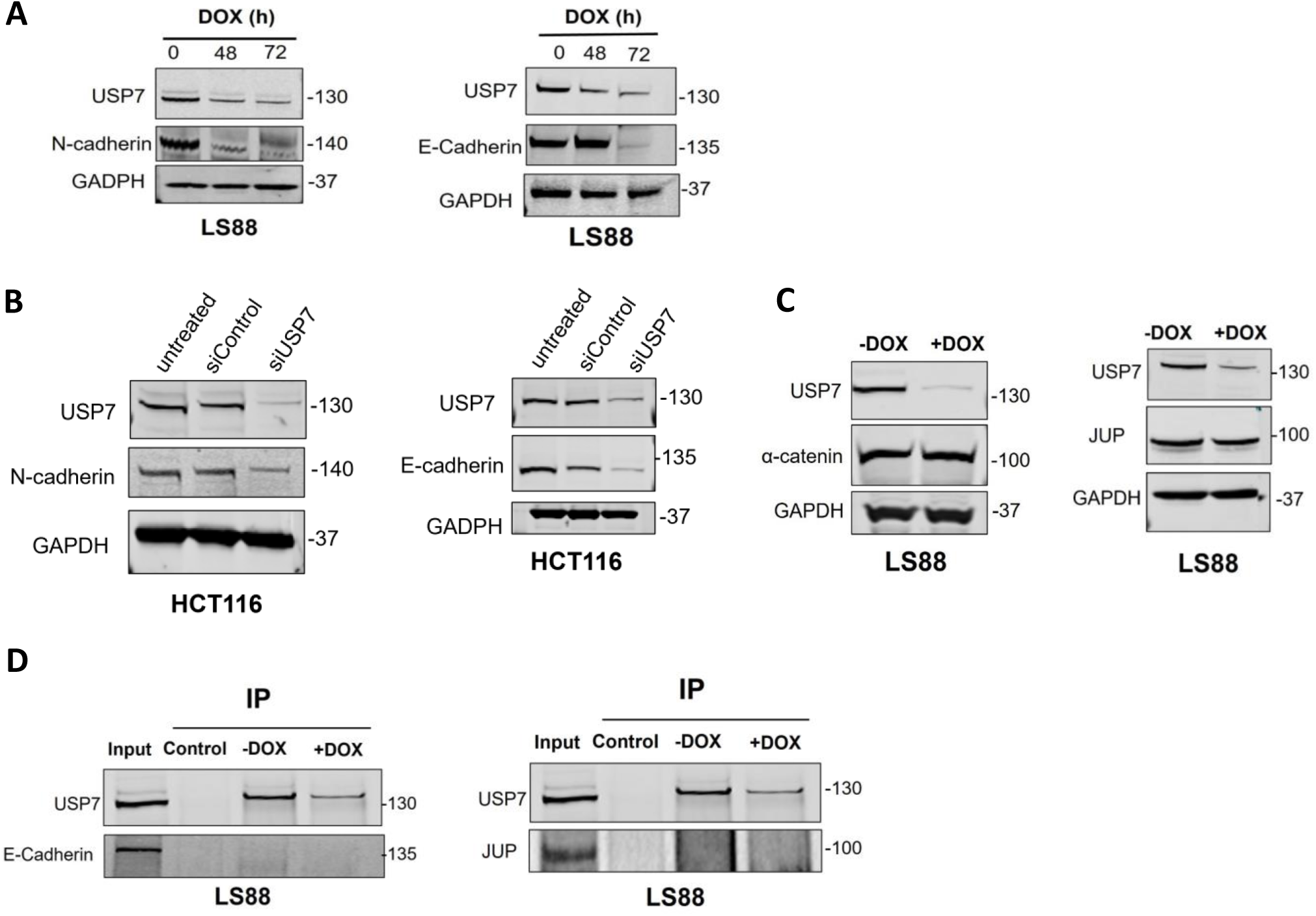
USP7 knockdown and effects on cadherin proteins and their substrates. **(A)** Protein expression of N-cadherin and E-cadherin in LS88 cells treated with doxycycline (DOX) for indicated time. GAPDH was used as a loading control. Adding doxycycline induces USP7 expression in LS88 cells. **(B)** Protein expression of N-cadherin and E-cadherin in HCT116 cells transfected with siUSP7 or siControl or left untreated. GAPDH was used as a loading control. **(C)** α-catenin and γ-catenin expression were analysed by western blot. N=3. GAPDH was used as a loading control. **(D)** Cell lysates from LS88 cells were immunoprecipitated with USP7, IgG antibodies followed by immunoblotting to examine the impact of USP7 knockdown on E-cadherin and γ-catenin (JUP).

Since Ajuba knockdown resulted in loss of cell-cell adhesion similarly to USP7 (Figure 6B), we next examined the expression of cell junction associated proteins in response to Ajuba knockdown. Since desmosomes and hemidesmosomes play key roles in cell-to-cell adhesion and cell-basement membrane adhesion (Green and Jones, 1996), we investigated the hemidesmosome protein integrin alpha 6 (ITGA6), the hemidesmosome protein integrin subunit beta 4 (ITGB4) and desmosomes protein Desmoglein-2 (DSG2) in doxycycline-treated LS88 cells and Ajuba knockdown cells. We observed a marked decrease in the expression of ITGB4 protein in Ajuba inhibited cells (Figure 8A-C), but no other detectable changes were observed.

**Figure 8.**
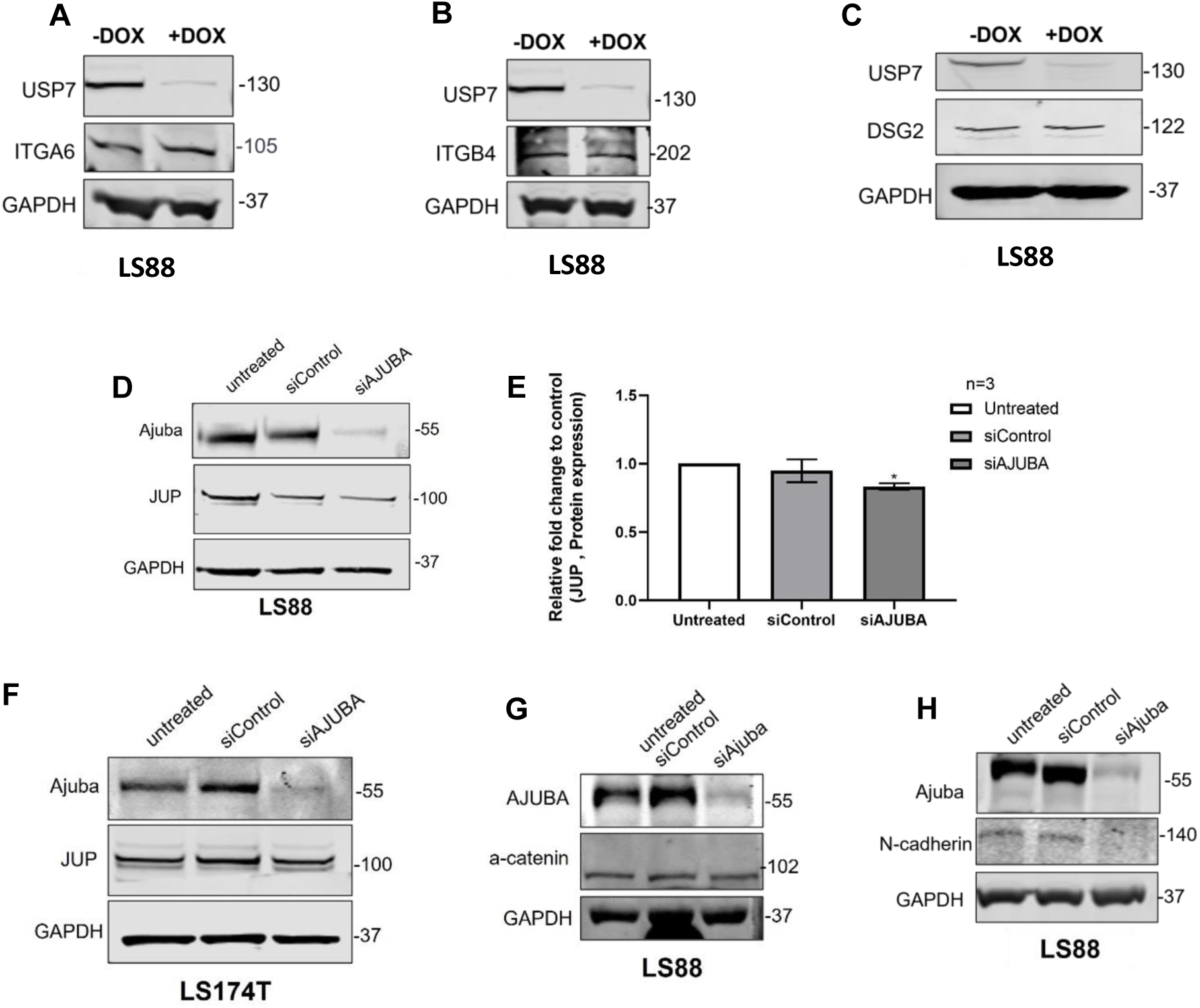
USP7 knockdown and effects on desmosome and hemidesmosome protein levels. **(A)** Protein expression of ITGA6 **(B)** ITGB4 and **(C)** DSG2 were analysed using western blot. GAPDH was used as a loading control. **(D)** LS88 cells were transfected with siControl and siAJUBA or left untreated, the cell lysates were subjected to western blotting with indicated antibodies and graph (E) showing protein levels of JUP in the three conditions. The Ordinary-One-way-ANOVA was used from three independent experiments. The significant difference between the two groups untreated and siAjuba groups is *P < 0.05. Mean ± SD. **(F)** Ajuba protein was knocked down in LS174T using siRNA and western blot was performed using AJUBA and JUP antibodies. **(G,H)** Cell lysates from LS88 cell were subjected to immunoblotting with AJUBA and α -catenin **and** N-cadherin antibodies. GAPDH was used as a loading control.

Next, LS88 cells were left untreated or transfected with siControl or siAJUBA, and a western blot was performed to examine JUP. We found a measurable decrease in JUP abundance in LS88 cells (Figure 8D and E), whereas no significant change was detected in JUP abundance in AJUBA depleted cells in LS174T (Figure 8F). We also examined α-catenin and N-cadherin protein levels in LS88. The most marked decreases in protein expression in Ajuba knockdown cells were observed for N-Cadherin (Figure 8G-H, consistent with previous reports (Marie et al., 2003; Sarpal et al., 2019).

Taken together, these analyses suggest that USP7 plays an indirect role in regulating cell-cell adhesion through its interaction and regulation of the LIM domain protein Ajuba.

## DISCUSSION

In this study, we used inducible knockdown proteomics to investigate novel interactions of the key deubiquitinase USP7 (Ubiquitin Specific Peptidase 7) in colorectal cancer. USP7 is known to promote cancer growth by upregulating several key cellular pathways, including the P53-MDM2 pathway (Harakandi et al., 2021) and Wnt/-catenin signaling (Novellasdemunt et al., 2017). Its role as a regulator of these key cancer pathways has led to USP7 becoming an important therapeutic target and yet our wider understanding of its substrates, interactions and functions is incomplete. We therefore used multi-modal approaches to investigate USP7 and identified the Ajuba LIM domain protein as a novel USP7 interaction partner.

We confirmed the substantial impact of USP7 on CRC cell viability in both 2D and 3D cultures, which is consistent with previous research that showed that USP7 inhibition using Parthenolide (PTL) reduced proliferation in HCT116 and SW480 CRC cells (Li et al., 2020).

The effect of USP7 depletion was further validated in relation to cell sphere volume in LS88, LS174T, and DLD-1. This revealed significant decreases compared to the HEK 293T cell line. In addition, our study has used colony formation assay to examine the role of USP7 in the regulation of cell colonies, thereby revealing substantial cell colony inhibition in depleted USP7 CRC cells. These findings concur with previous research (Behan et al., 2019; Hayal et al., 2020). The current study verifies the importance of USP7 for cell survival in cases of colorectal cancer.

Cell adhesion plays an essential role in metastasis and cancer progression. The interaction between cancer cells and the endothelium defines the metastatic spread (Bendas and Borsig, 2012). One pathway that was enriched in our MS data for USP7 is cell adhesion. To characterize the USP7 role in cell-cell junction, USP7 was knocked down, and the impacts on cell-cell adhesion were observed using dispase-based assay and mechanical stress in LS88. The results showed significant reductions in cell junctions. In the subsequent stage, we examined the master proteins that play a major role in cell adhesion in USP7-depleted cells in our cell model LS88 and other CRC cells. The findings confirmed the low expression of N-cadherin and E-cadherin when USP7 levels are reduced, although transcription levels must be measured in this context. However, no physical interactions were found between USP7, N-cadherin, E-cadherin or JUP.

Many studies have shown that Ajuba acts as an oncogene due to its regulatory role in key signaling pathways, such as RAS/ERK, JAK/STAT, Wnt and Hippo. The protein acts as a co-regulator of main transcription factors, e.g. Sp1, Snail and nuclear hormone receptors. Amplified Ajuba expression has been found in viral types of tumours, suggesting its potential as a clinical prognostic and diagnostic marker (Jia et al., 2020). Ajuba is one of the proteins that has been linked to cell adhesion for over a decade (Marie et al., 2003; Nola et al., 2011). The bioinformatics results emerging from our MS data indicated that Ajuba is a novel substrate for USP7 in our cell model. These findings were acquired in the lab using immunochemistry, immunoprecipitation, and siRNA oligos. The data confirmed the MS data, thereby indicating that Ajuba interacts directly with USP7 in CRC cells. We subsequently tested the central hypothesis that USP7 may influence cell-cell adhesion by regulating Ajuba protein levels. The results showed a reduction of N-cadherin and ITGB4 protein levels in Ajuba downregulated cells. These findings are consistent with earlier research (Peifer and Wleschaus, 1990). The results from the dispase-based essay in our lab demonstrated the high number of fragments in Ajuba knocked down cells. When combined, these findings suggest that the inhibition in the cell-cell junction in USP7-depleted cells may occur via Ajuba protein levels.

## DATA AVAILABILITY

The mass-spectrometry data has been submitted to the PRIDE database with the following details: Project accession: PXD039488. All other data is provided within this paper and in the supplementary data.

## ACKNOWLEDGEMENTS

RME acknowledges Medical Research Council (MR/S01411X/1) and European Commission FP7 (“Oncoprotnet”) for funding; YW acknowledges Medical Research Council (MR/S025480/1). The authors thank Professor Madelon Maurice (University Medical Center Utrecht) for kindly providing us with the LS88 cell-line.

## SUPPLEMENTARY DATA

**Supplementary Data 1**

Supplementary Figures

**Supplementary Table 1**

Full results from Mascot search engine analysis of all mass-spectrometry data

**Supplementary Table 2**

Normalized area quantifications from Proteome Discoverer analysis of mass-spectrometry data

## Notes

**Conflict of Interest Disclosure**: The authors declare that there are no known conflicts of interest

### Competing Interest Statement

The authors have declared no competing interest.

