## Supplementary Data 1 for "Knockdown proteomics reveals USP7 as a regulator of cell-cell adhesion in colorectal cancer via AJUBA"

Supplemental Data

### TITLE

Knock-down proteomics reveals HAUSP/USP7 as a regulator of cell-cell adhesion in colorectal cancer via the LIM domain protein AJUBA

### AUTHORS AND AFFILIATIONS

Ahood Al-Eidan^1,2^, Ben Draper^1^, Siyuan Wang^1^, Brandon Coke^1^, Paul Skipp^1^, Yihua Wang^1^, Rob M. Ewing^1*^

1 School of Biological Sciences, Faculty of Environmental and

Life Sciences, University of Southampton, Southampton,

United Kingdom

2 Department of Biology, College of Sciences, Imam

Abdulrahman Bin Faisal University, P.O. Box 1982, Dammam, Saudi Arabia.

*

Postal address of submitting author: Life Sciences Building 85,

Highfield campus, University of Southampton, Southampton, HANTS, SO17 1BJ, UK

**A**

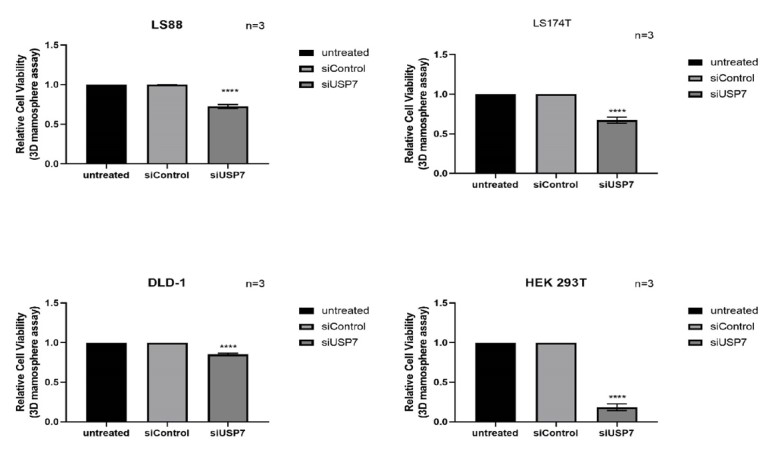

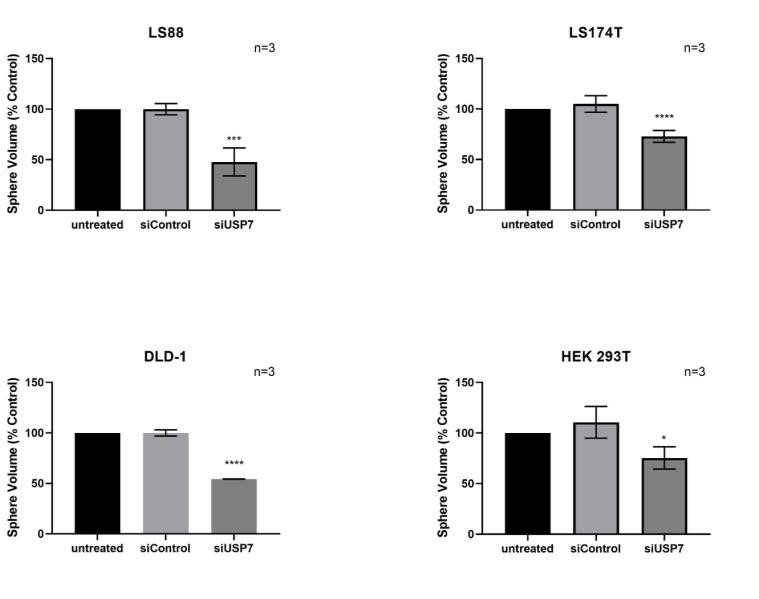

**B**

**C**

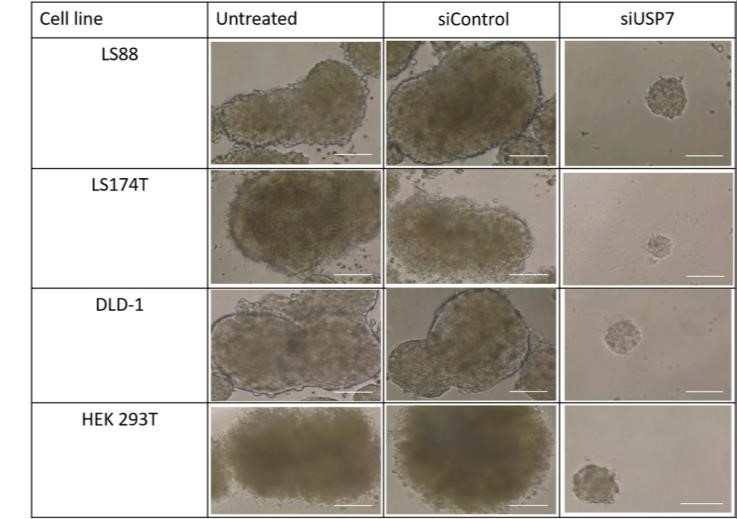

**Fig. S1: USP7 is required for the survival of colorectal cancer cells cultured in 3D.**

**(A)** Graph showing relative cell viability in (untreated and transfected with siControl or USP7 siRNA) colorectal cell lines in 3D cultures using Cell-Titer Glo® assay. **(B)**, Graphs showing sphere volume for colorectal cancer cell lines and HEK 293T cells after transfections after two weeks cultured in 3D. **(C),** 3D images for LS88, LS174T, DLD-1 and HEK 293T cell lines. Scale bar: 250 µm. For statistical analysis, Ordinary one-way ANOVA was performed. Data are mean ± SD . n= three independent experiments. The significant difference between the two groups untreated and siUSP7 groups is (*P < 0.05. **P < 0.01. ***P <

0.001. ****P < 0.0001).

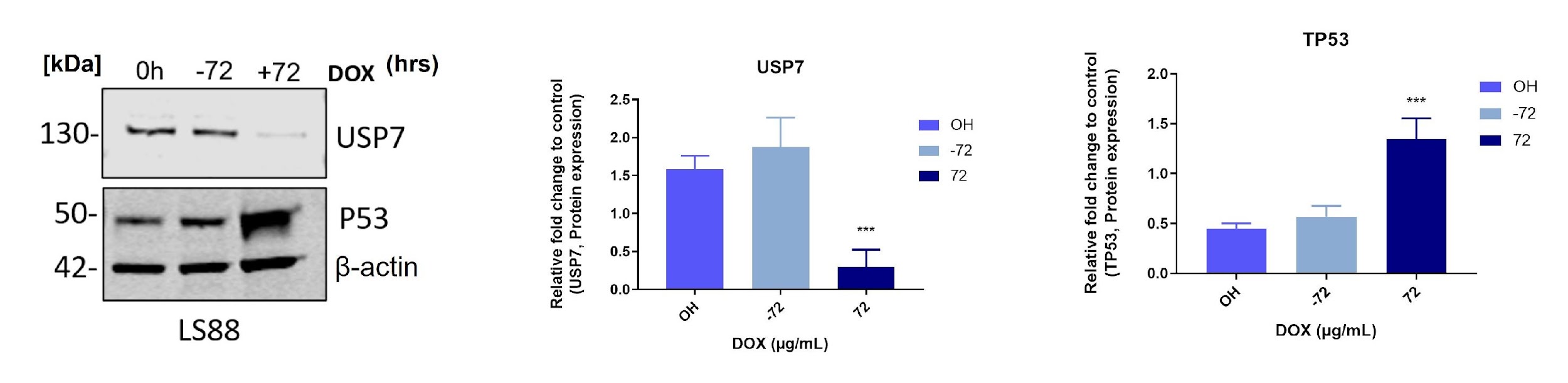

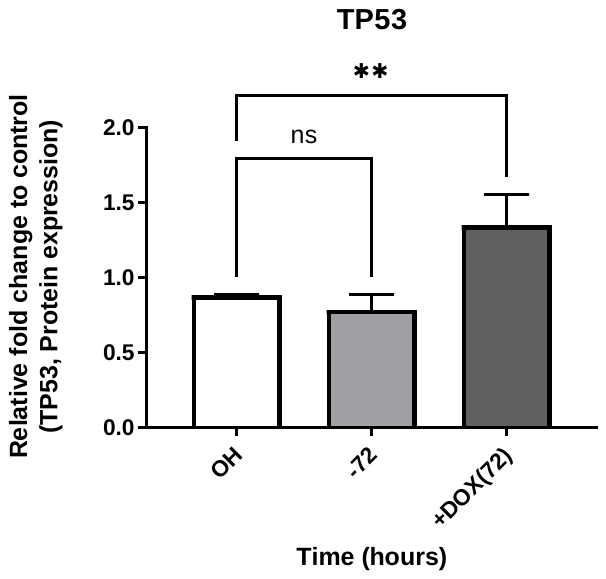

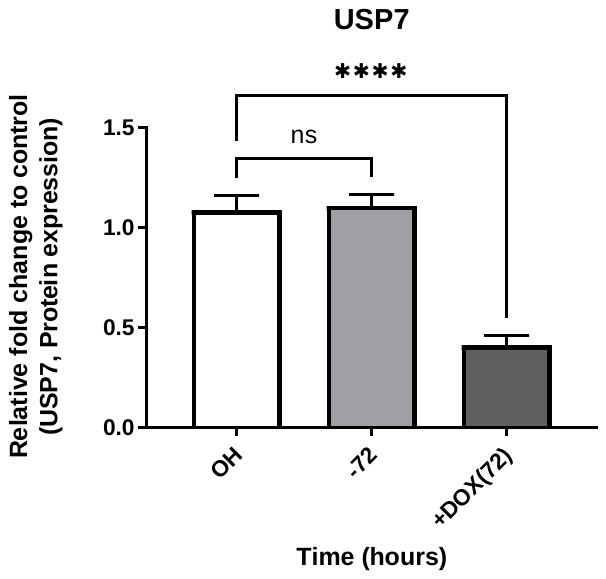

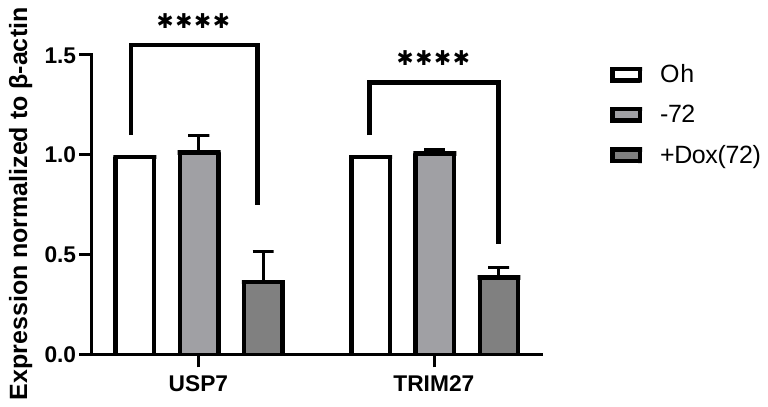

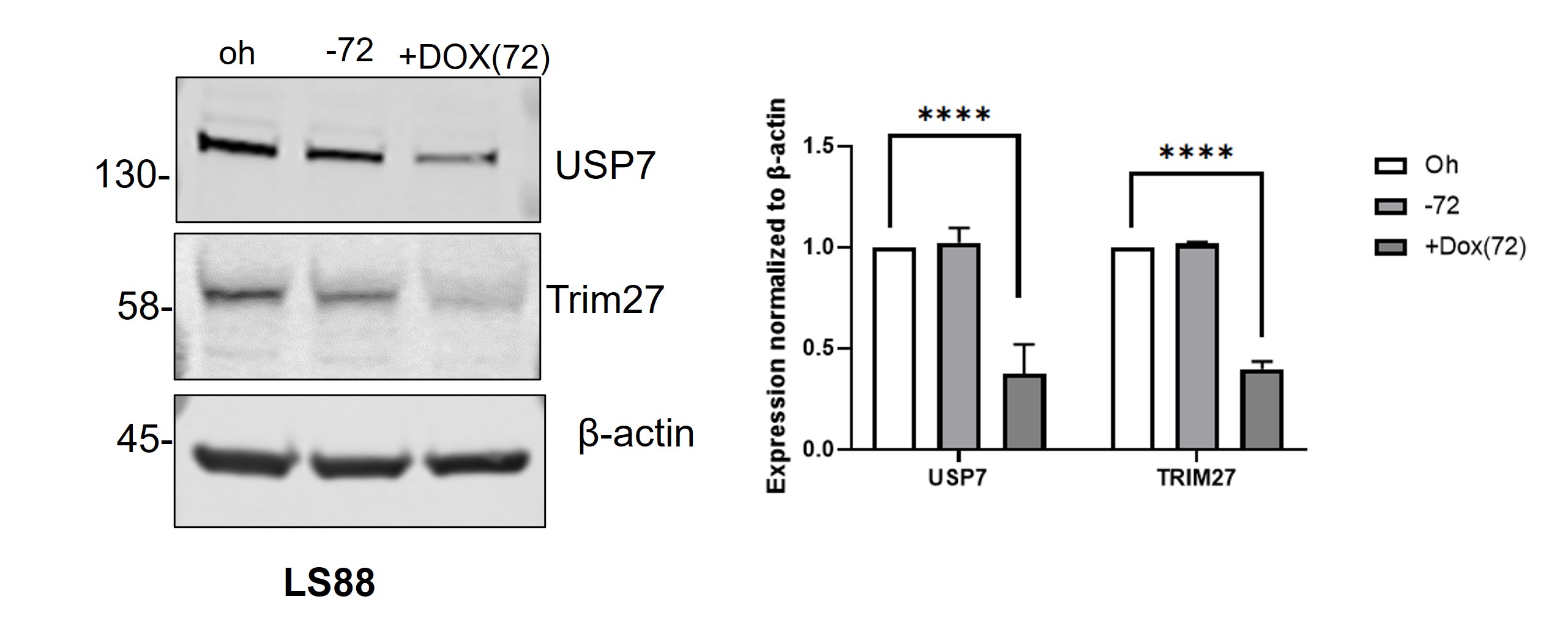
A

B

**Fig. S2: USP7 regulates P53 and Trim27 in LS180 colorectal cancer cells**

**(A)** Immunoblotting analysis for USP7 and P53 in LS88. The protein levels for USP7a and TP53 after knocked down USP7 in LS88. One-way ANOVA was performed for statistical analysis. Data presented as the mean ± the SD.

**(B)** LS88 cells were treated for the specified time points with 1 µg/mL doxycycline and control cells were analysed using immunoblotting for protein expression of Trim27. The Ordinary-One-way-ANOVA Dunnett's multiple comparisons test was performed, the mean ± SD determined from three independent experiments. (***p < 0.001) and (****P ≤ 0.0001).

**A**

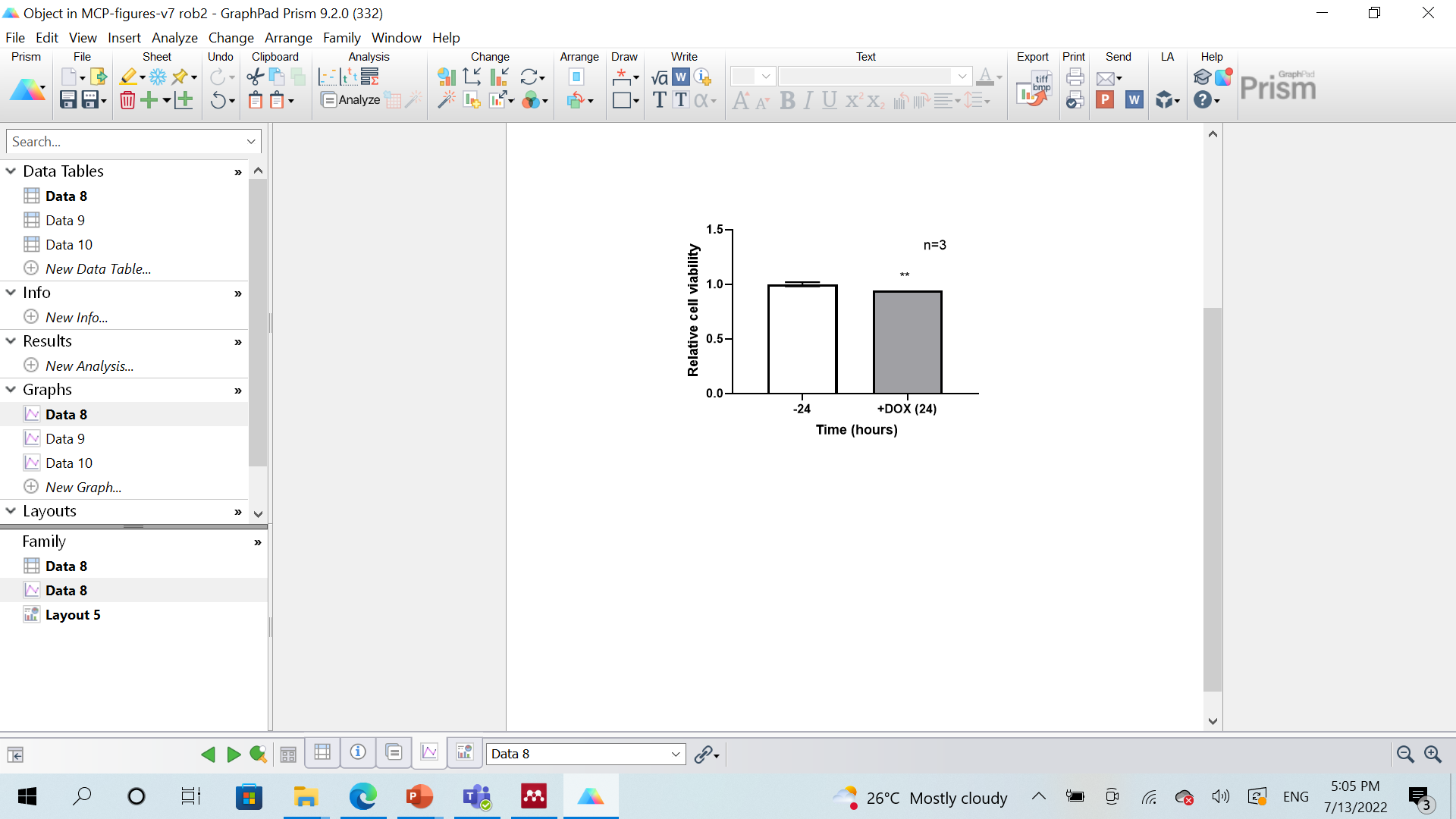

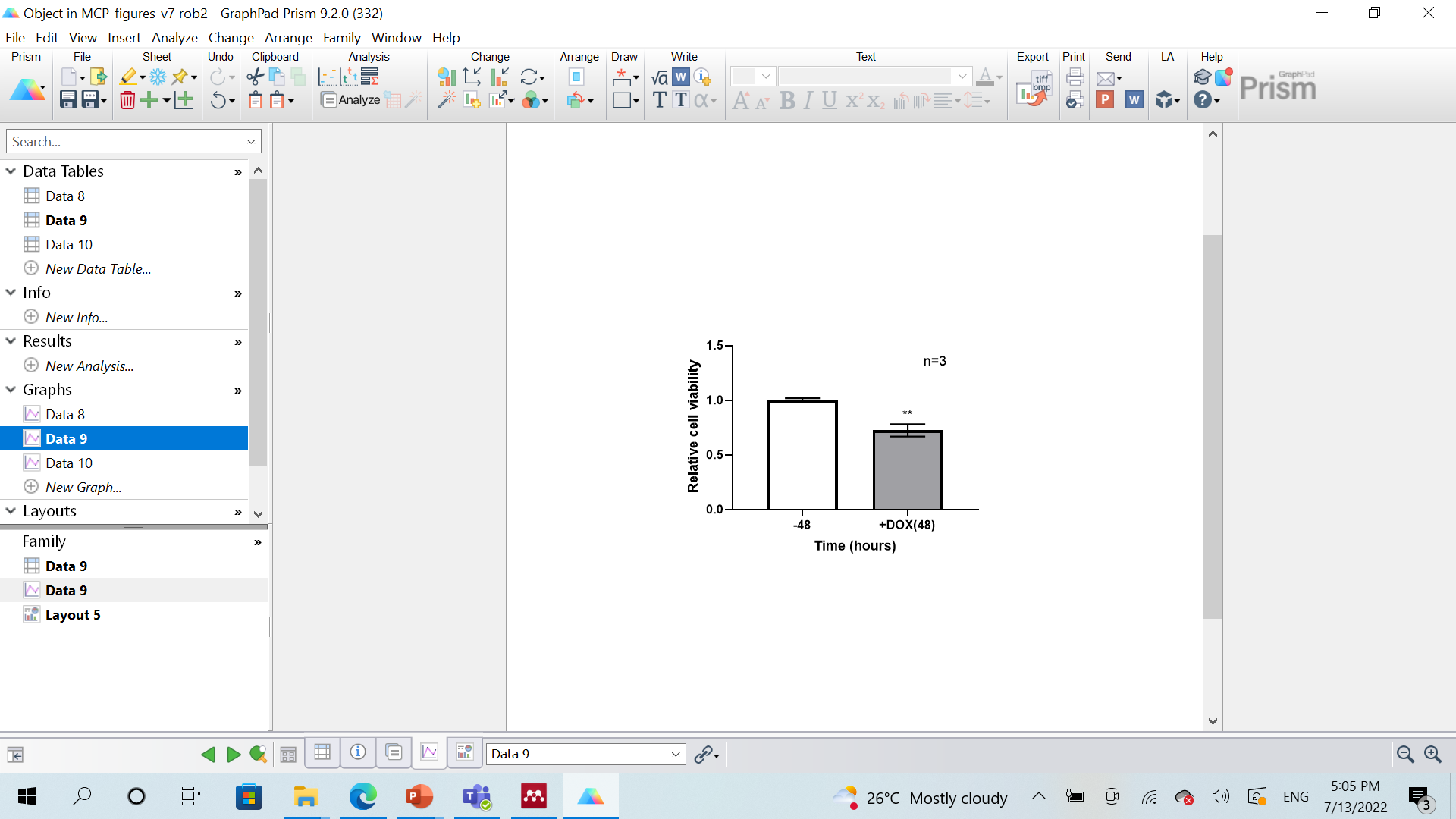

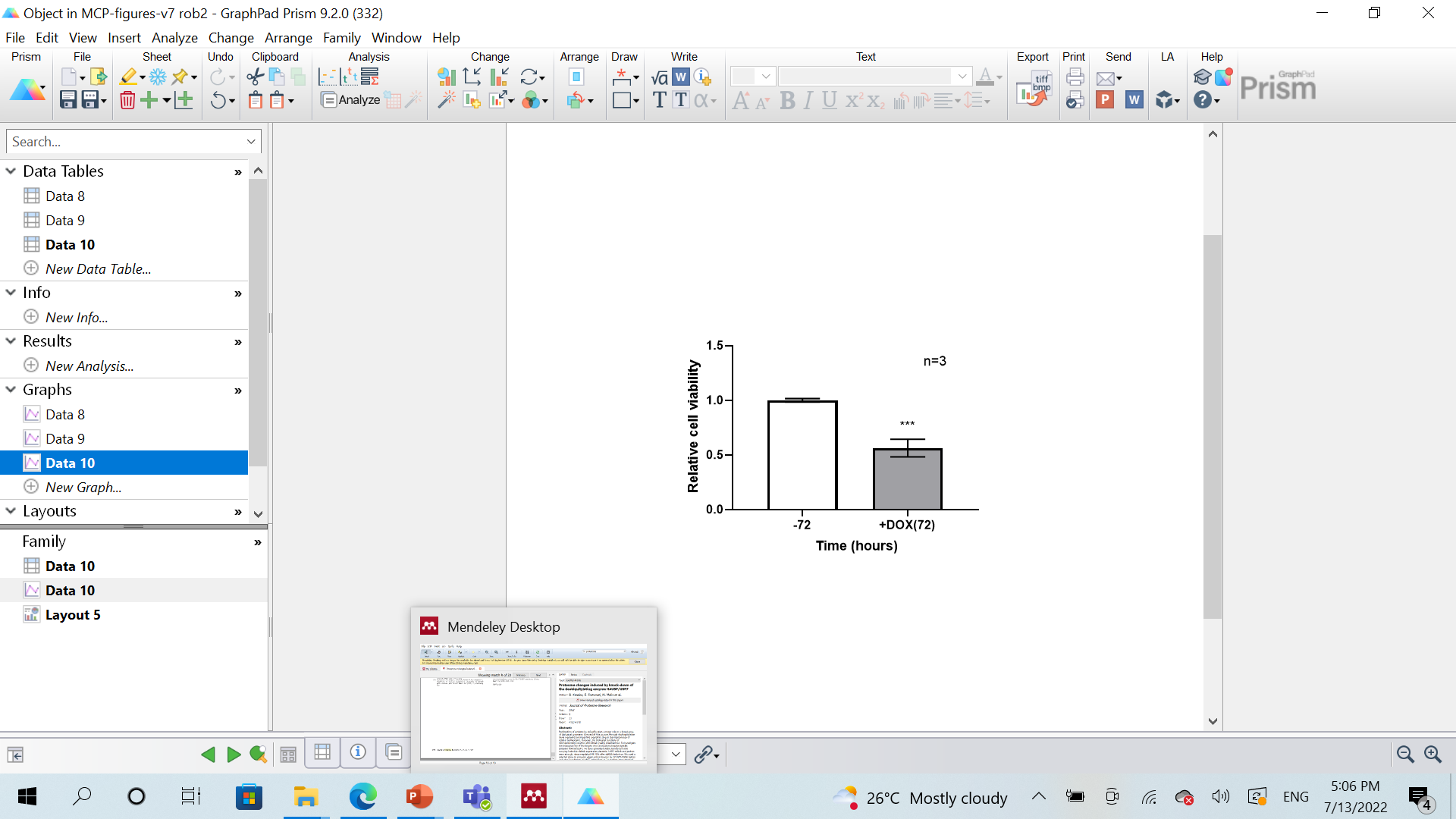

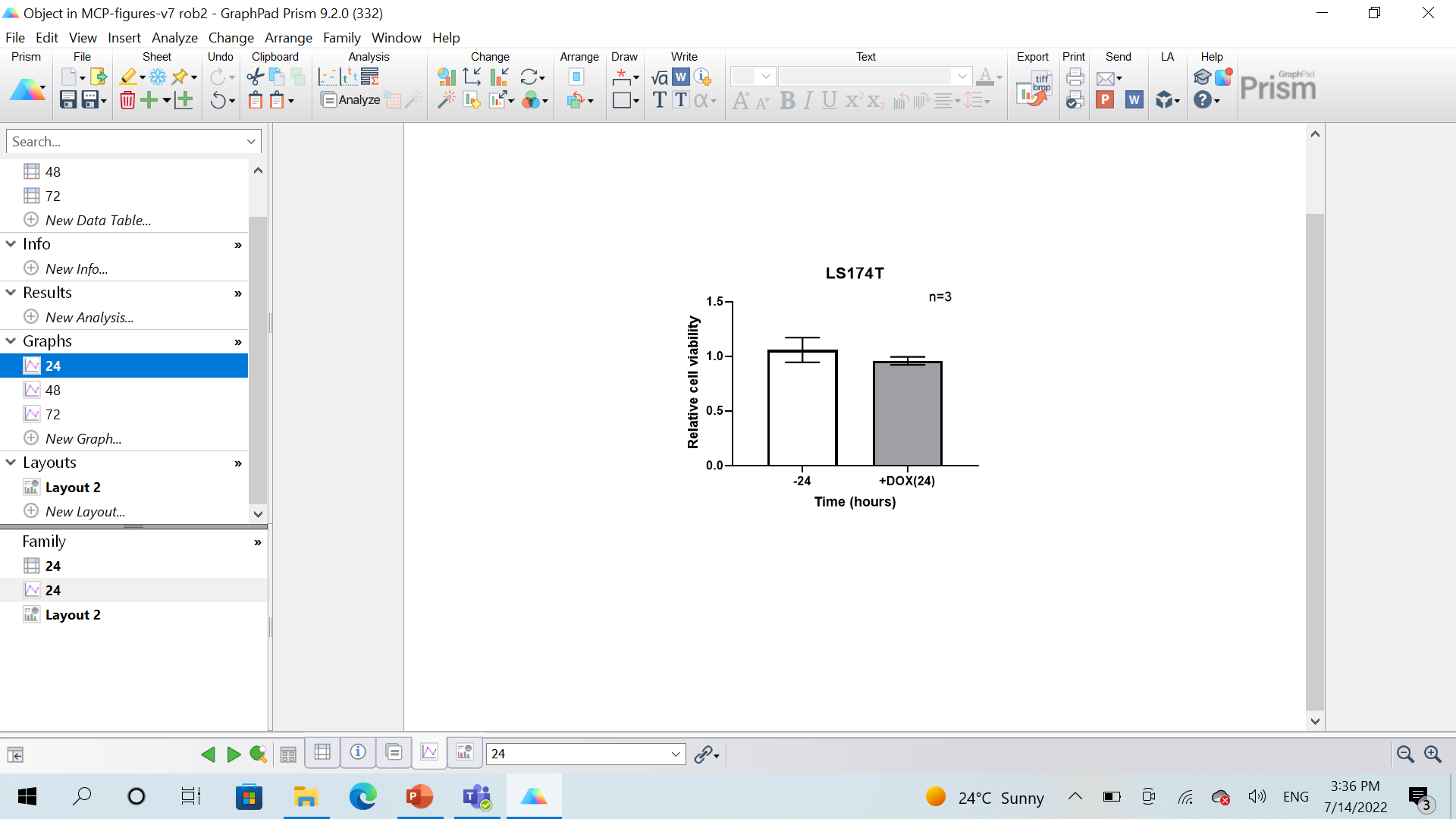

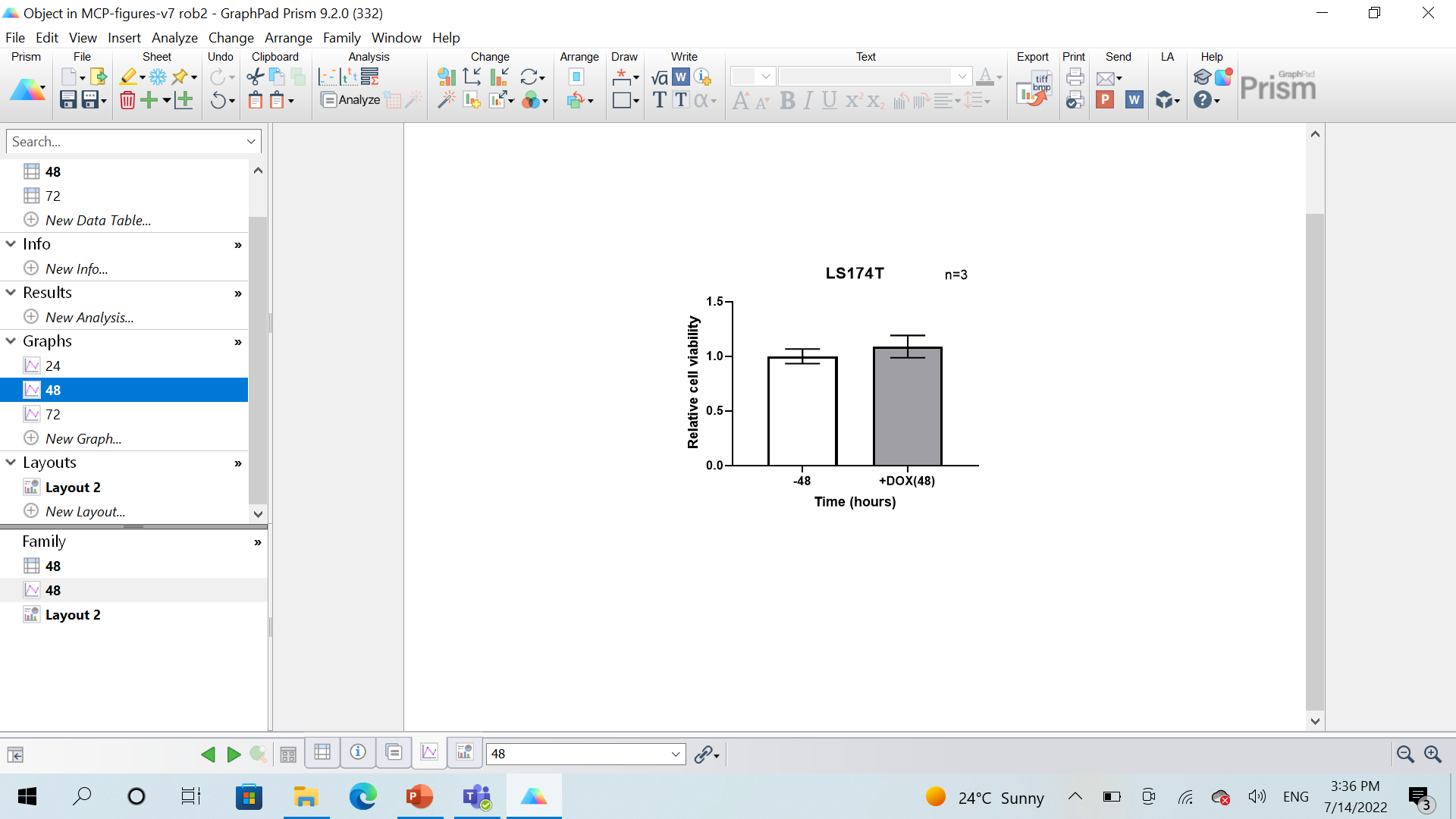

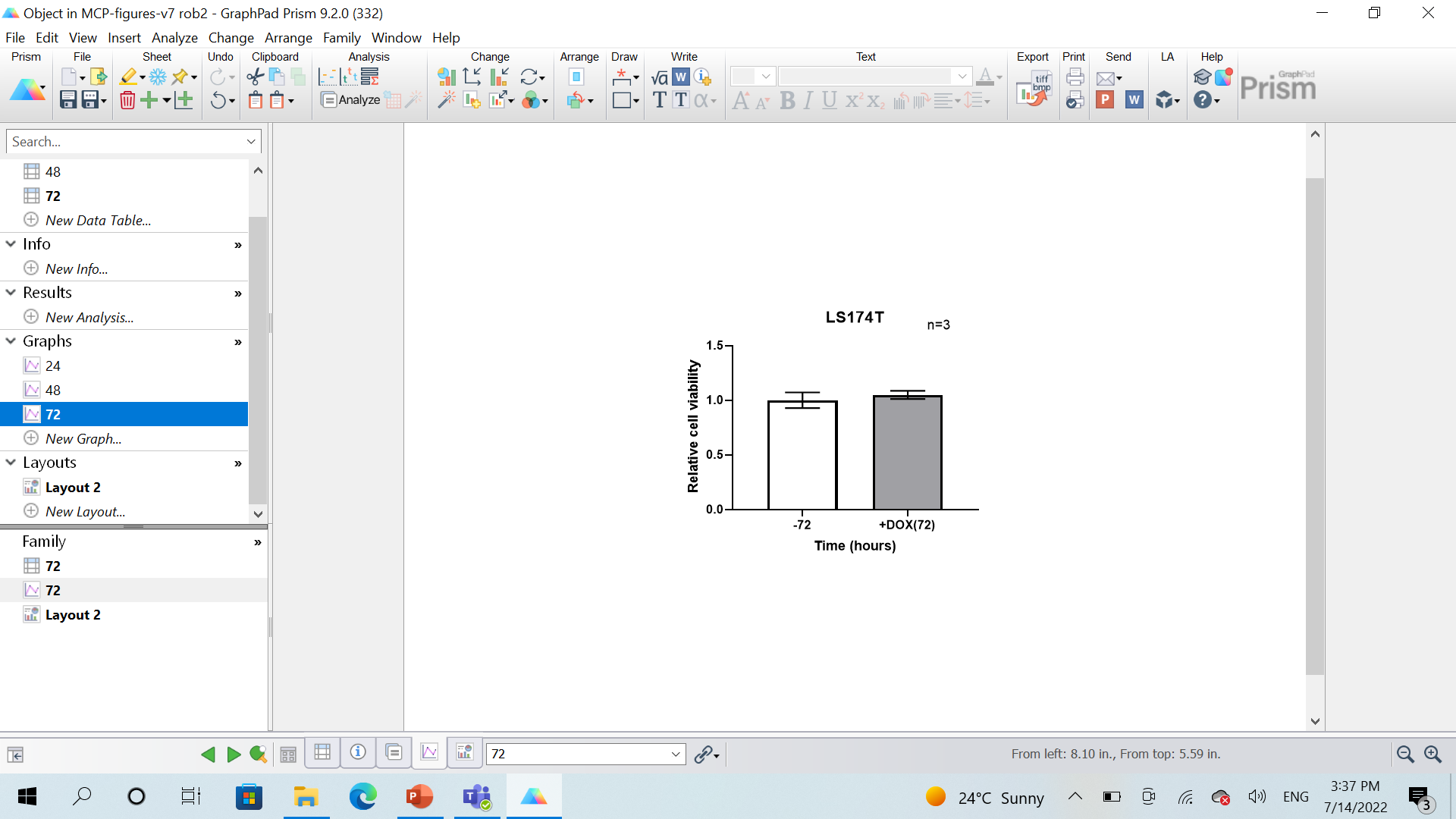

B

C

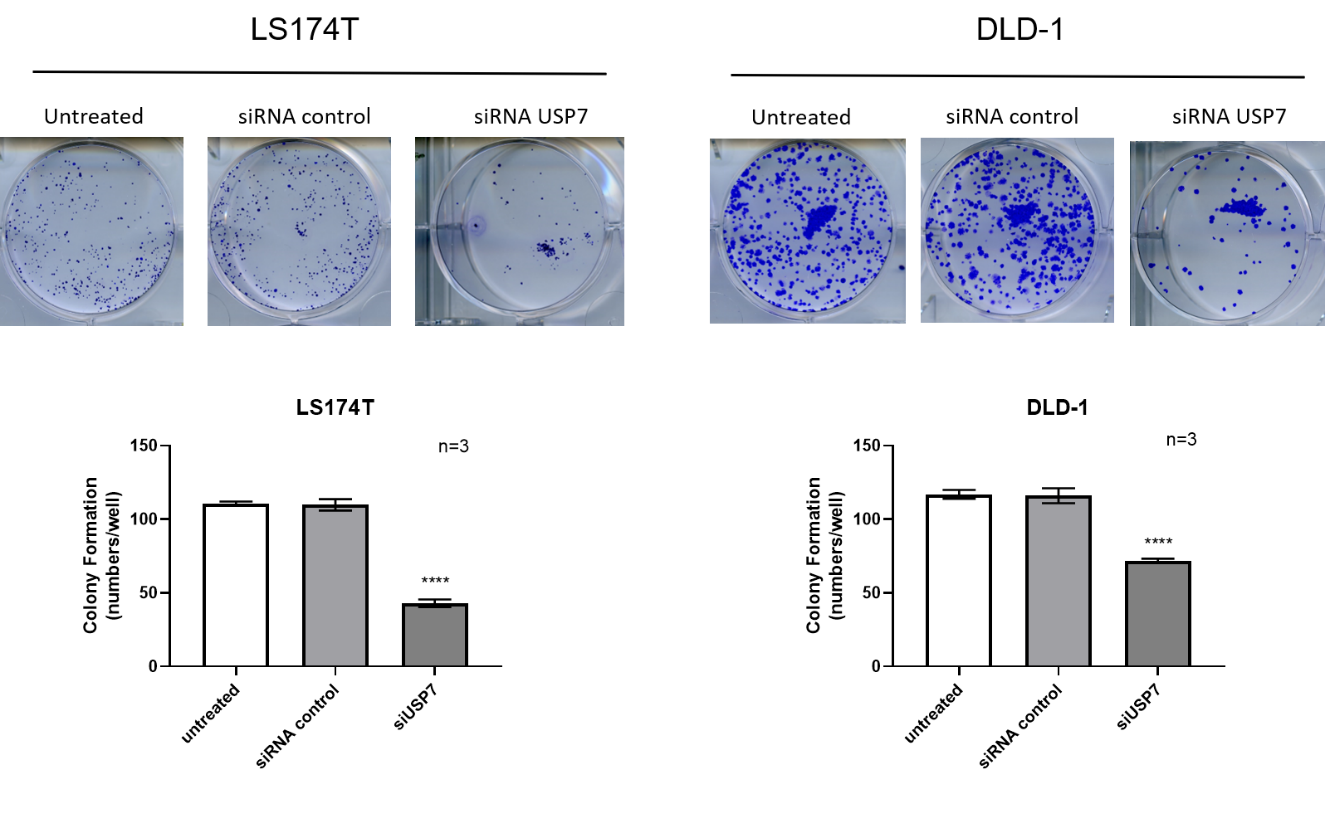

**Fig. S3: Effects of USP7 depletion on cell viability and clonogenicity of CRC cells**

Cells were treated for the specified time points with 1 µg/mL concentrations of doxycycline over a period of 72 hours. The number of viable cells was assessed using a CellTitre assay (Promega). (**A)** in LS88 which contain inducible shRNA targeting USP7. ** = significant difference between untreated and treated cells after 24 and 48 hours, p-value <0.01. *** = significant difference between untreated and treated cells after 72 hours, p-value <0.001 (Student’s t-test; two tailed unpaired), mean ± SD. (**B)** in LS174t cells**. (C)** LS174T and DLD-1 were transfected with USP7 siRNA before plated for colony formation assay. Untreated cells served as a reference control in the assay. Data are presented as mean ± SD (one way-a nova) from at least three independent experiments; P ≤ 0.0001.

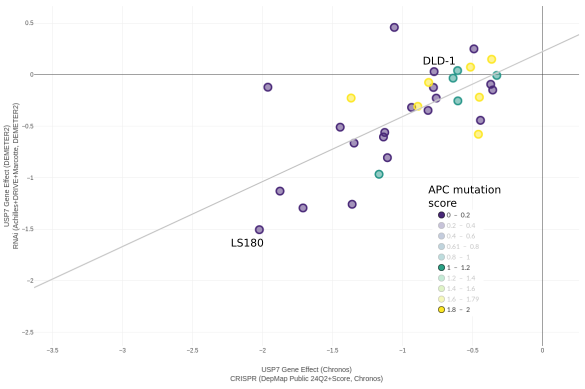

**Fig. S4: Dependency Map (depmap.org) analysis of USP7 and β-catenin expression in colorectal cancer cell lines.**

Scatterplot comparing the distribution of USP7 gene effect processed scores (CERES, DEMETER2) between the RNAi dataset (y) and CRISPR dataset (x). All available colorectal cancer cell-lines are included and are coloured according to APC mutation hotspot score. The plot indicates a marked separation of USP7 gene effect according to APC mutation.

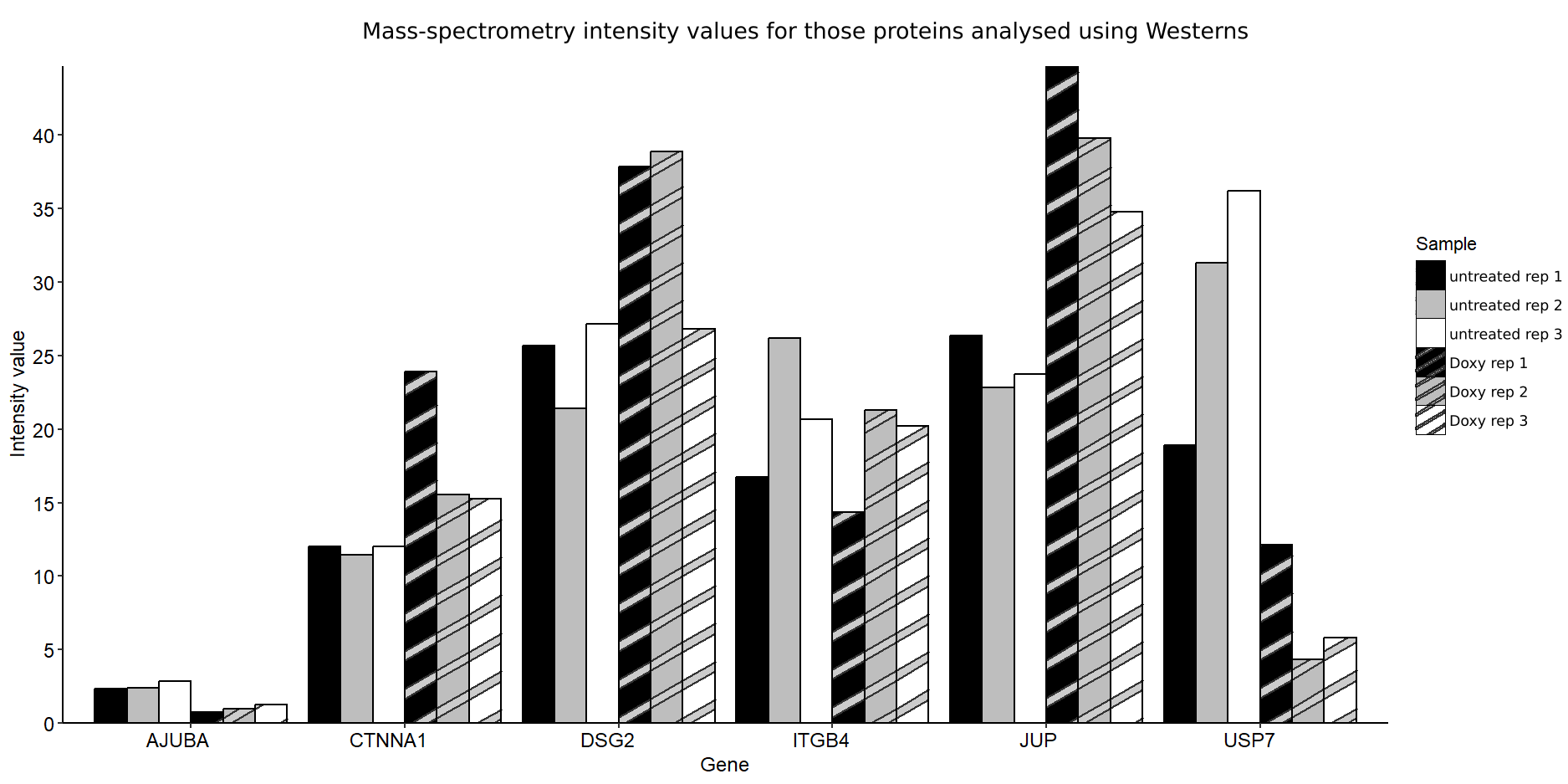

**Fig. S5: Protein intensity values from the Mass-spectrometry analysis for those proteins analysed using Westerns**

Protein intensity values from the mass-spectrometry data comparing untreated and treated (Doxycycline-inducible USP7 knockdown) samples.

**A**

**FT671**

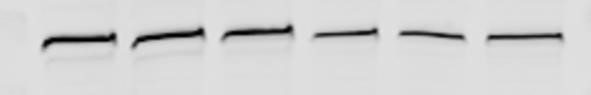

USP7

0

h

h

24

72

h

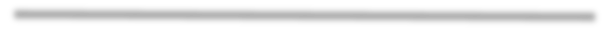

10

μ

M

4

h

48

h

96

h

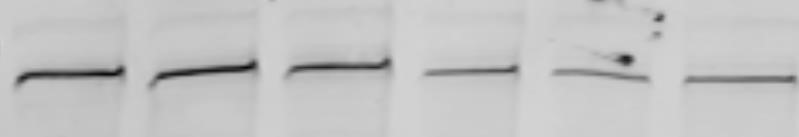

N

-

cadherin

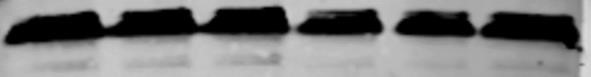

GAPDH

-

130

-

37

-

140

**HCT116**

# B FT671

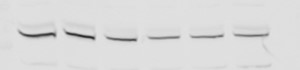

0

h

24

h

72

h

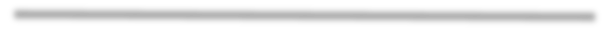

10

μ

M

4

h

48

h

96

h

USP7

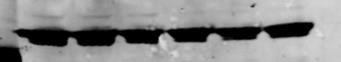

GAPDH

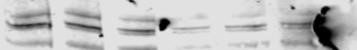

N

-

cadherin

-

130

-

37

-

140

**LS88**

**Fig. S6: FT671 downregulates N-cadherin expression in colorectal cancer cells.**

FT671 treatment dose-dependently decreased levels of N-cadherin in **(A)** Hct116 **(B)** LS88.

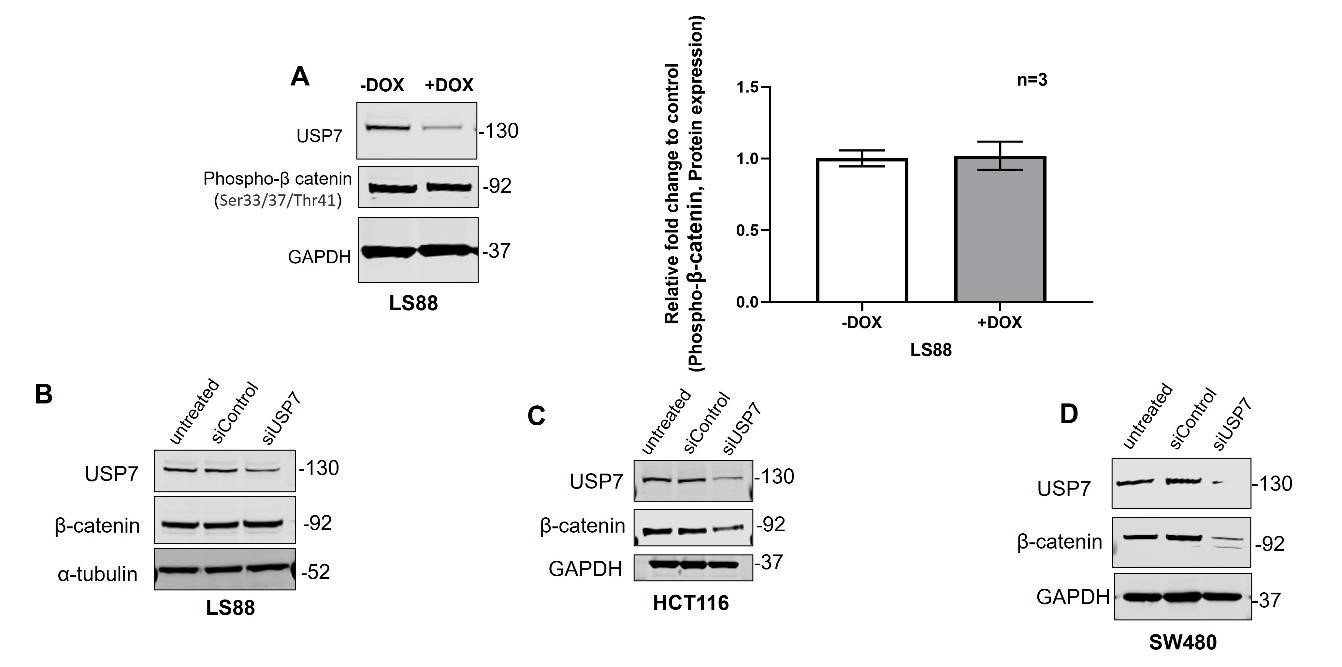

**Fig. S7: USP7 and β-catenin expression in colorectal cancer cell lines.**

**(A)**& **(B)** LS188 (CTNNB1 S45F/S45F), **(C)** HCT116 (CTNNB1 S45del/WT) and **(D)** SW480 (WT/WT). The immunoblotting shows showing reduced levels of B-catenin in HCT116. GAPDH and α-Tubulin were used as a control. The mean value ± SD was obtained from three independent experiments. Two tailed unpaired t- test was determined to identify statistical differences.

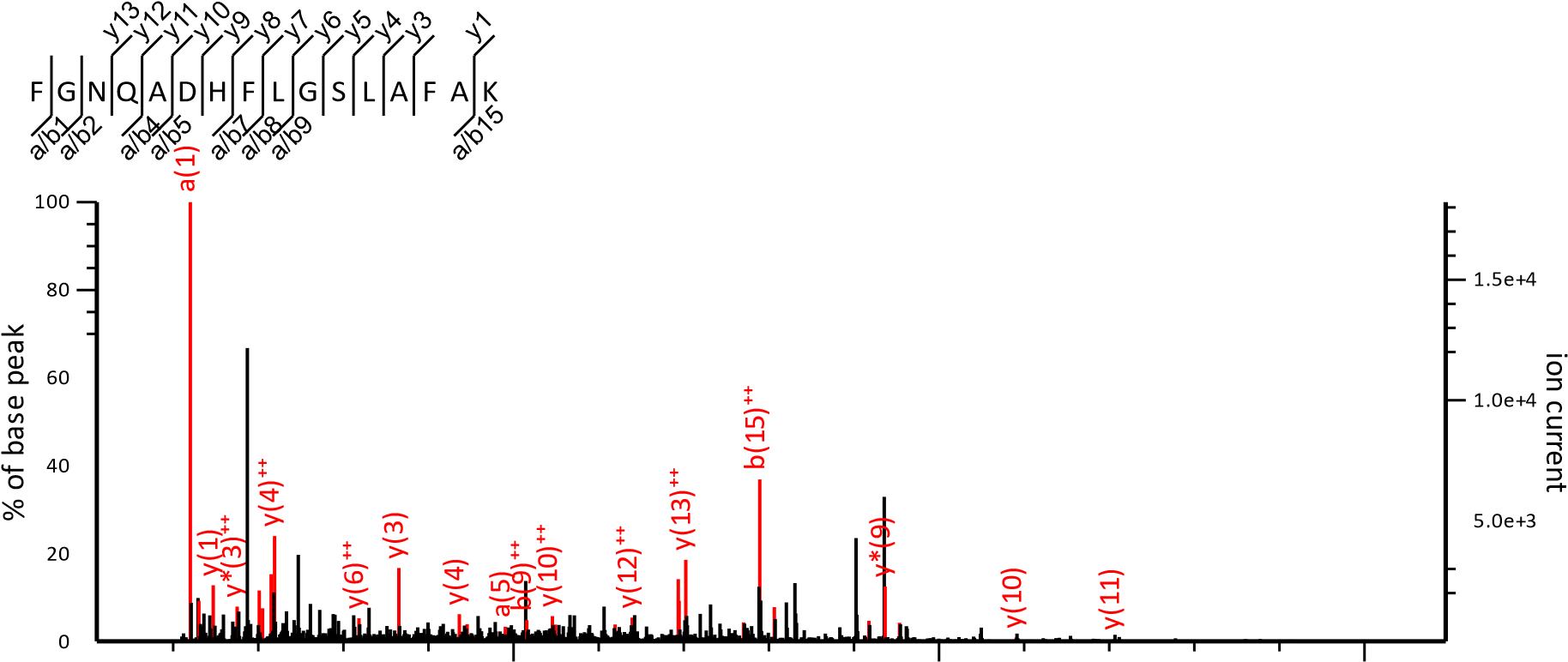

1	1

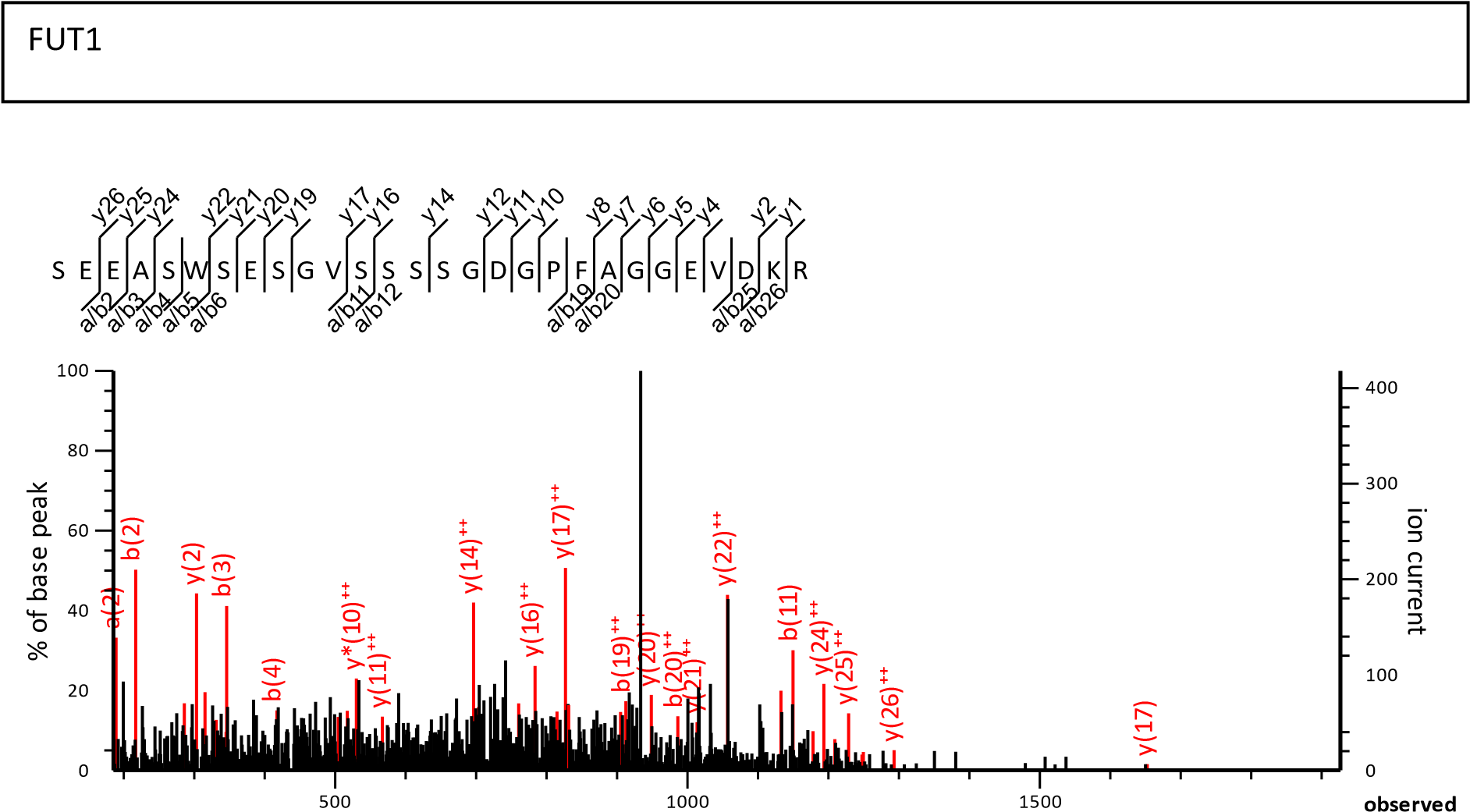

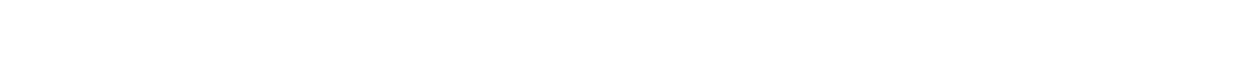

CC1

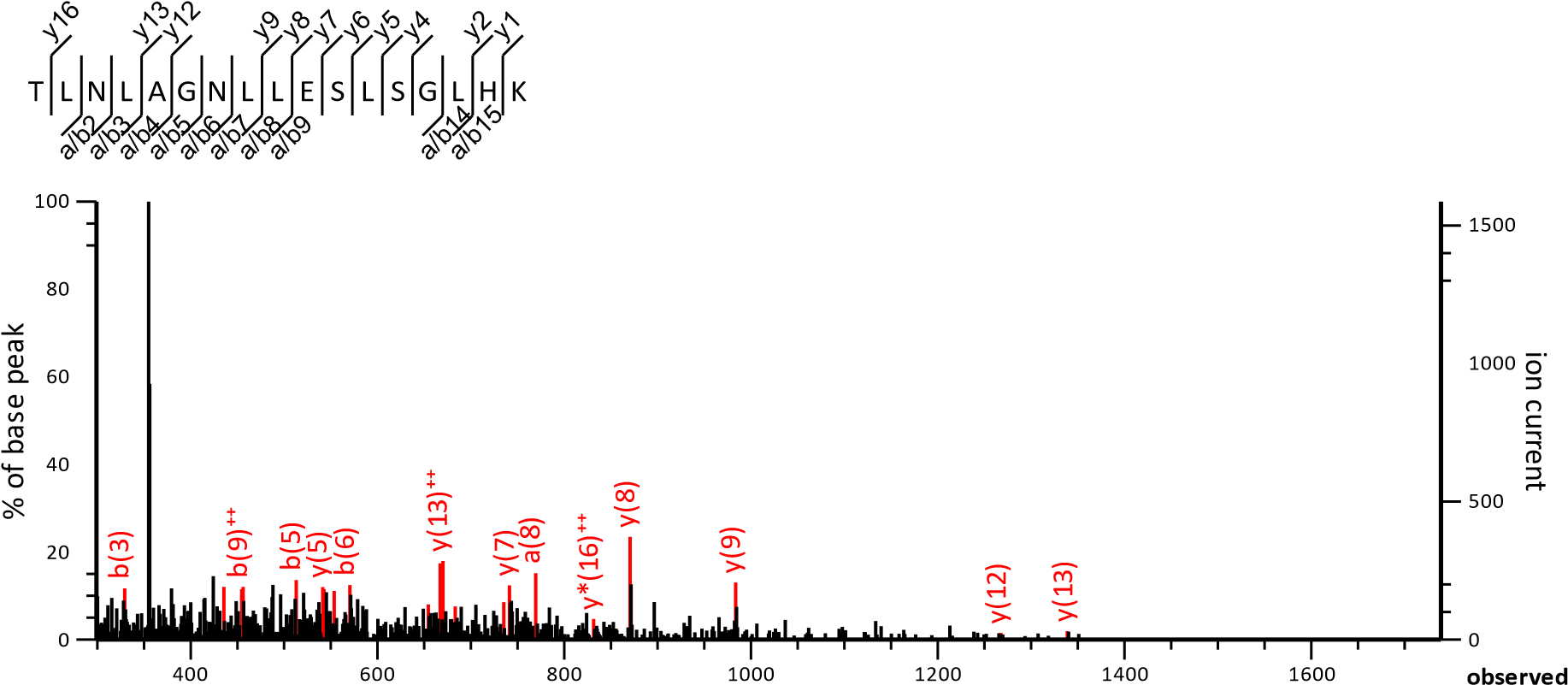

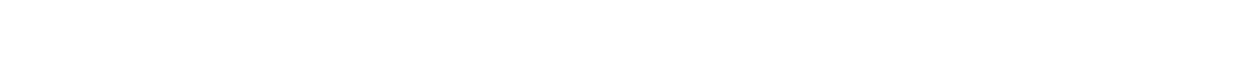

I C

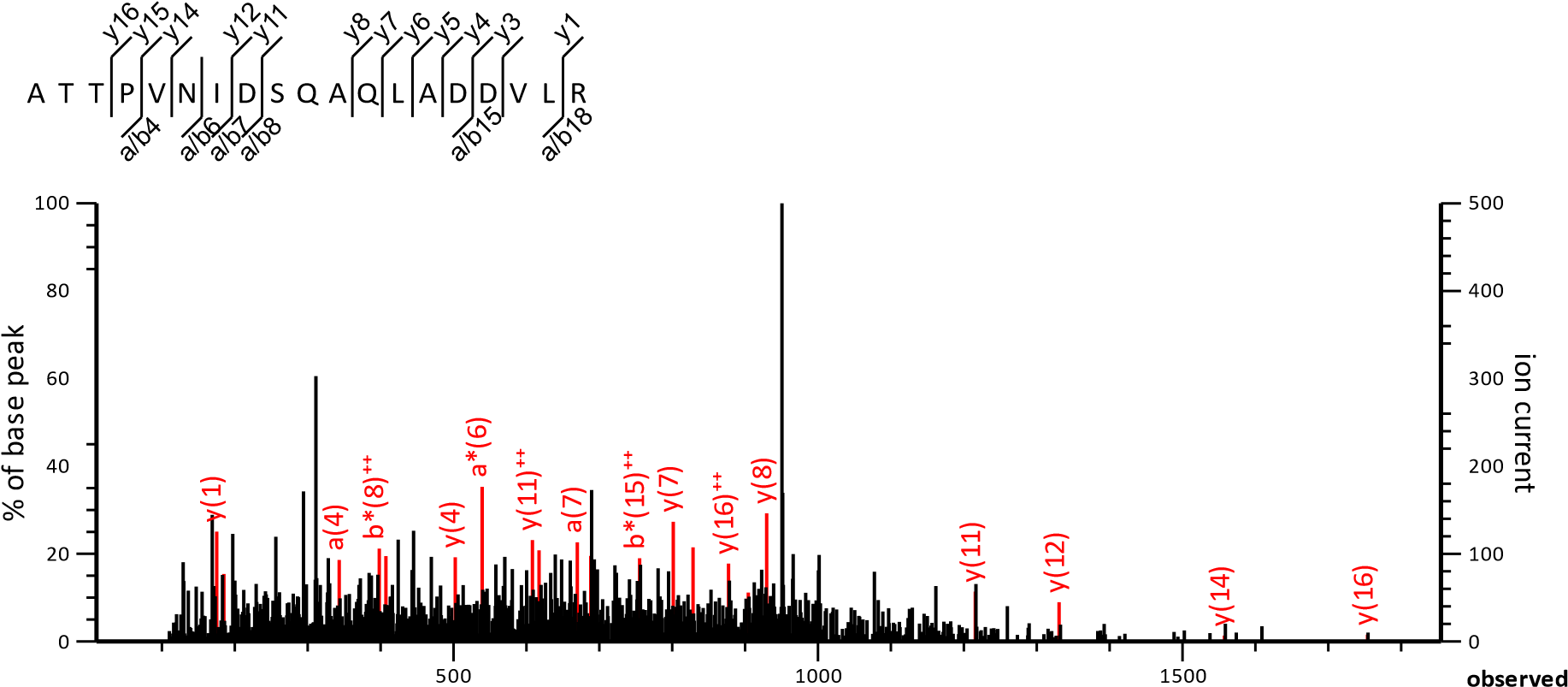

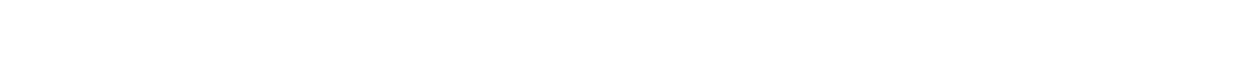

1

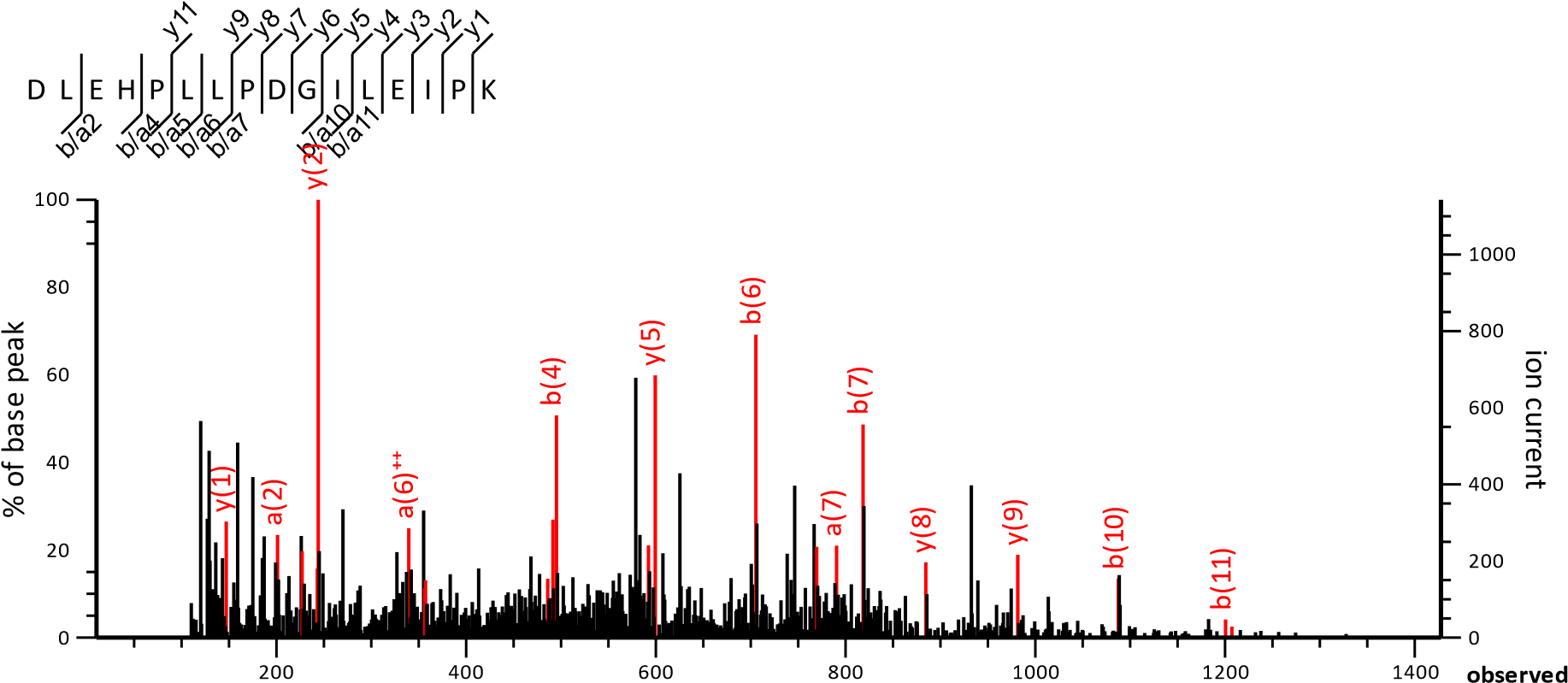

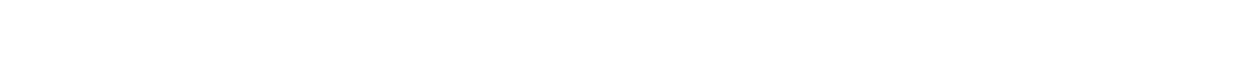

MI1

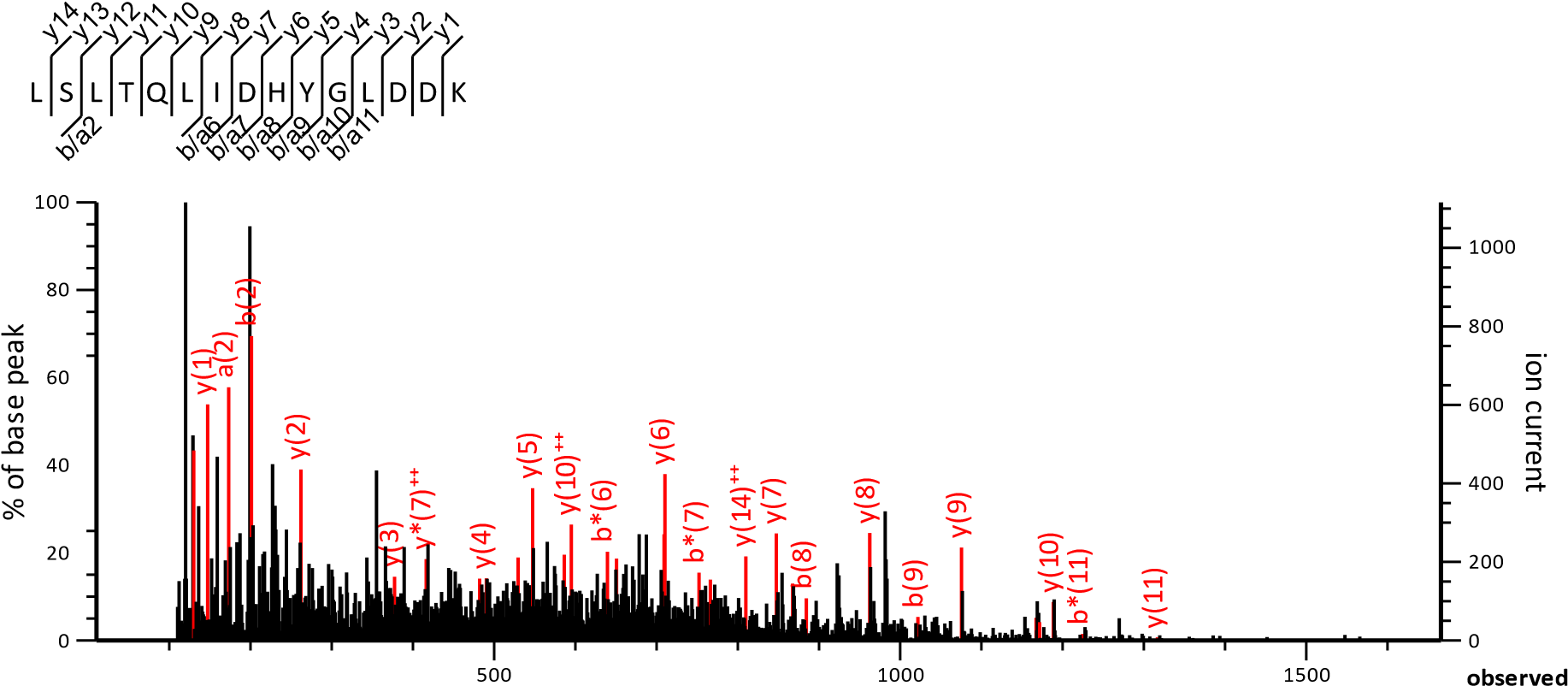

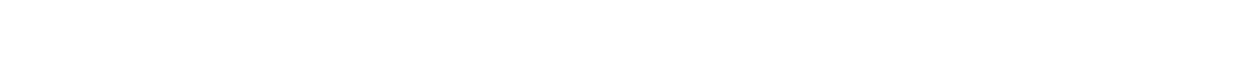

F 1

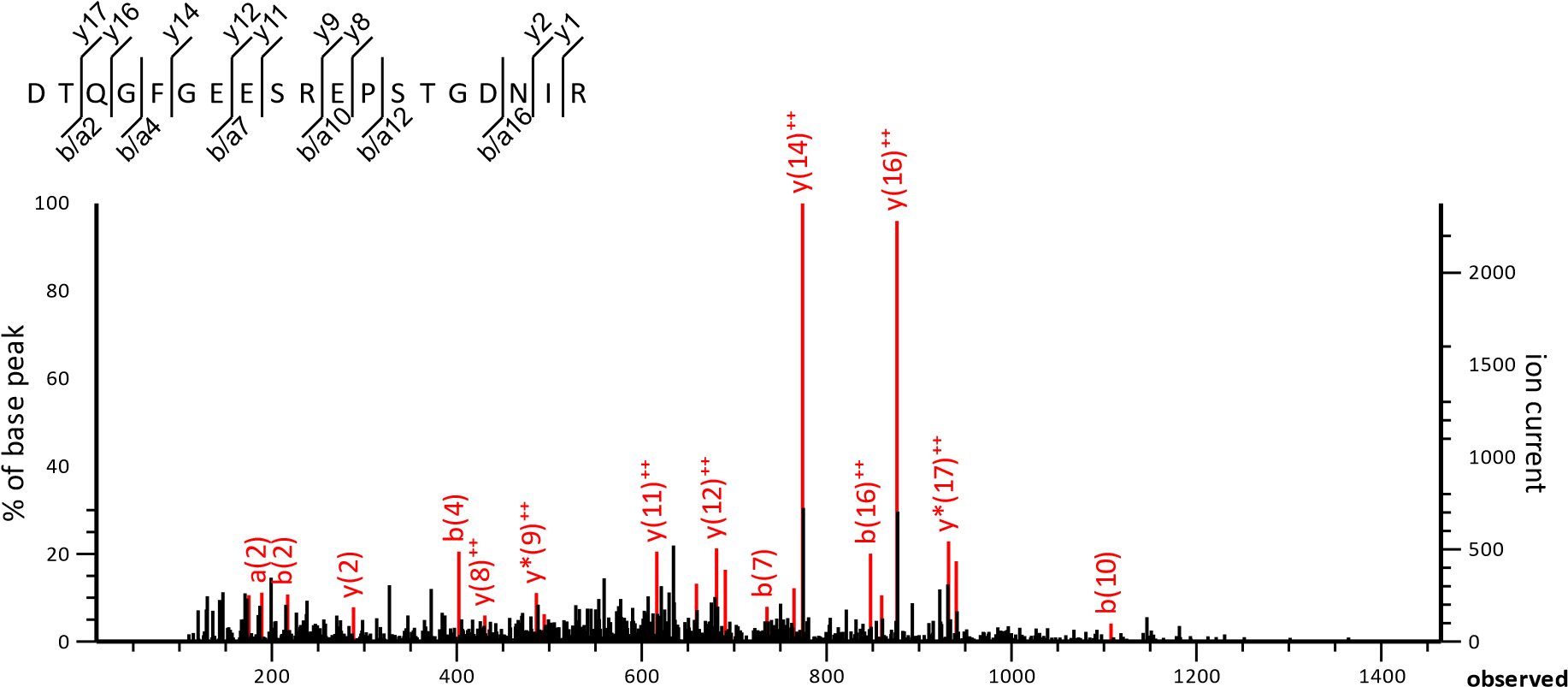

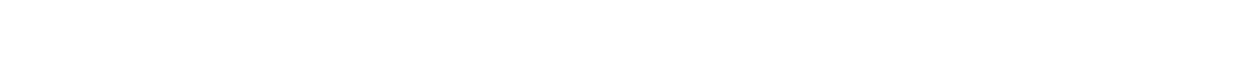

MIM

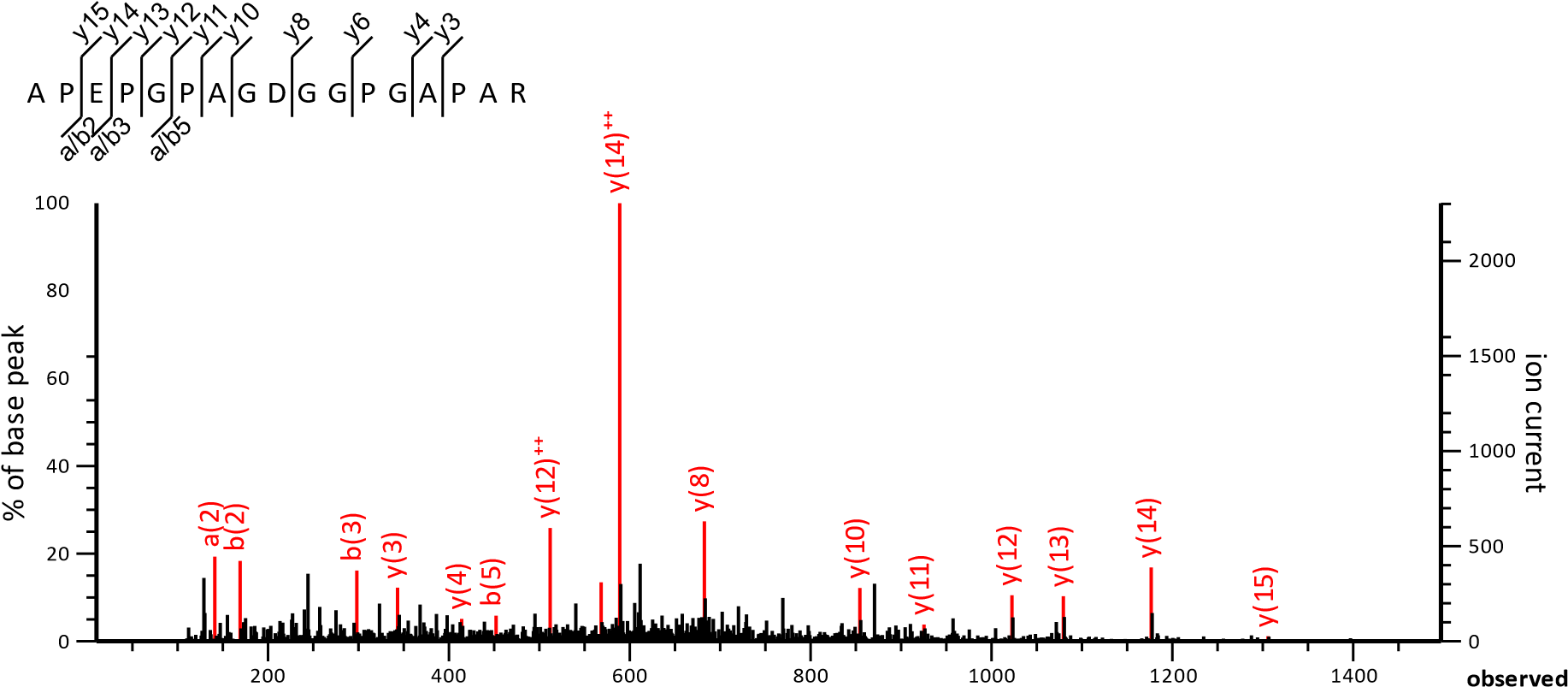

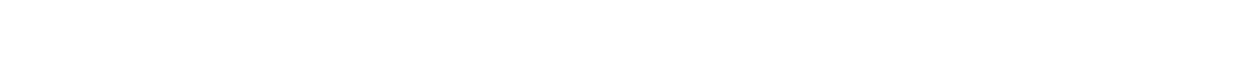

X1

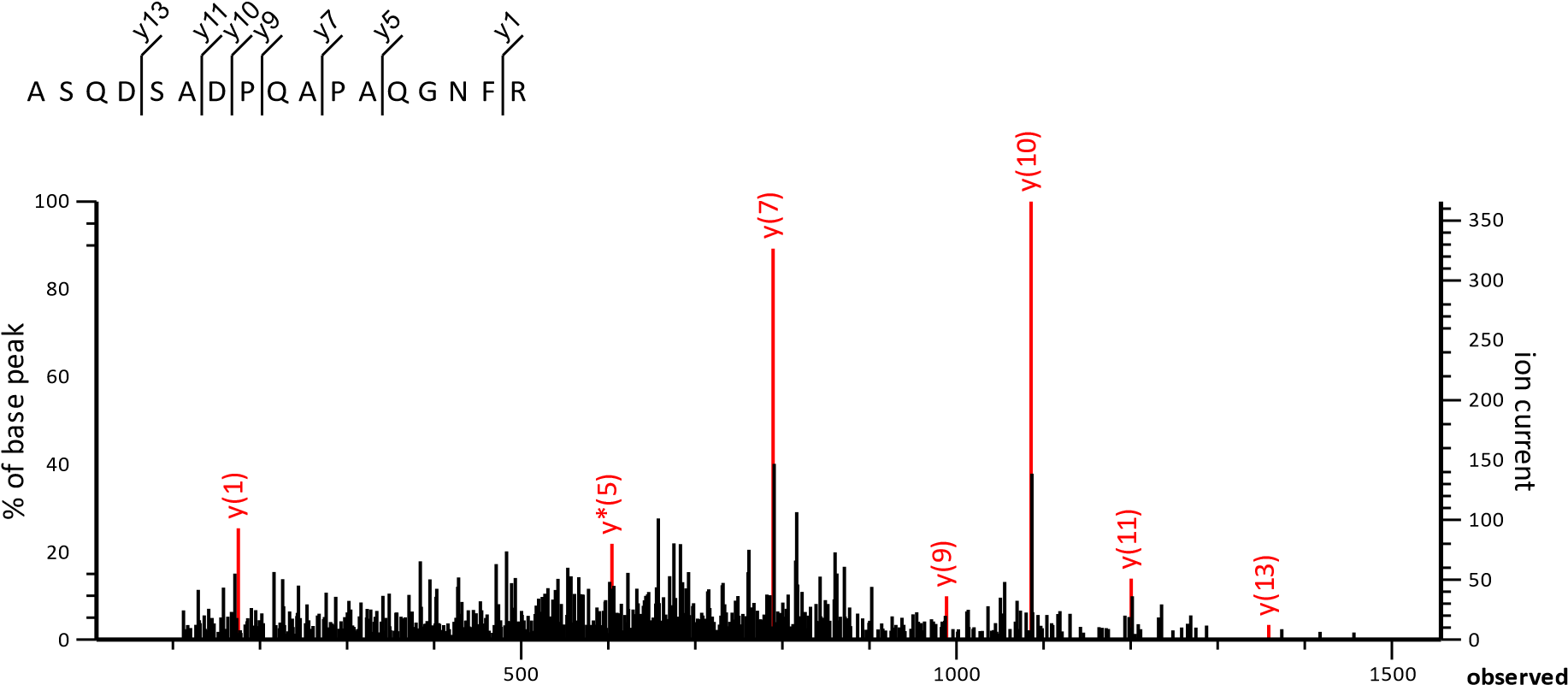

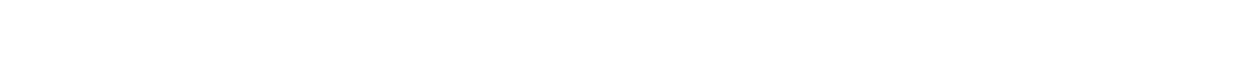

OX1

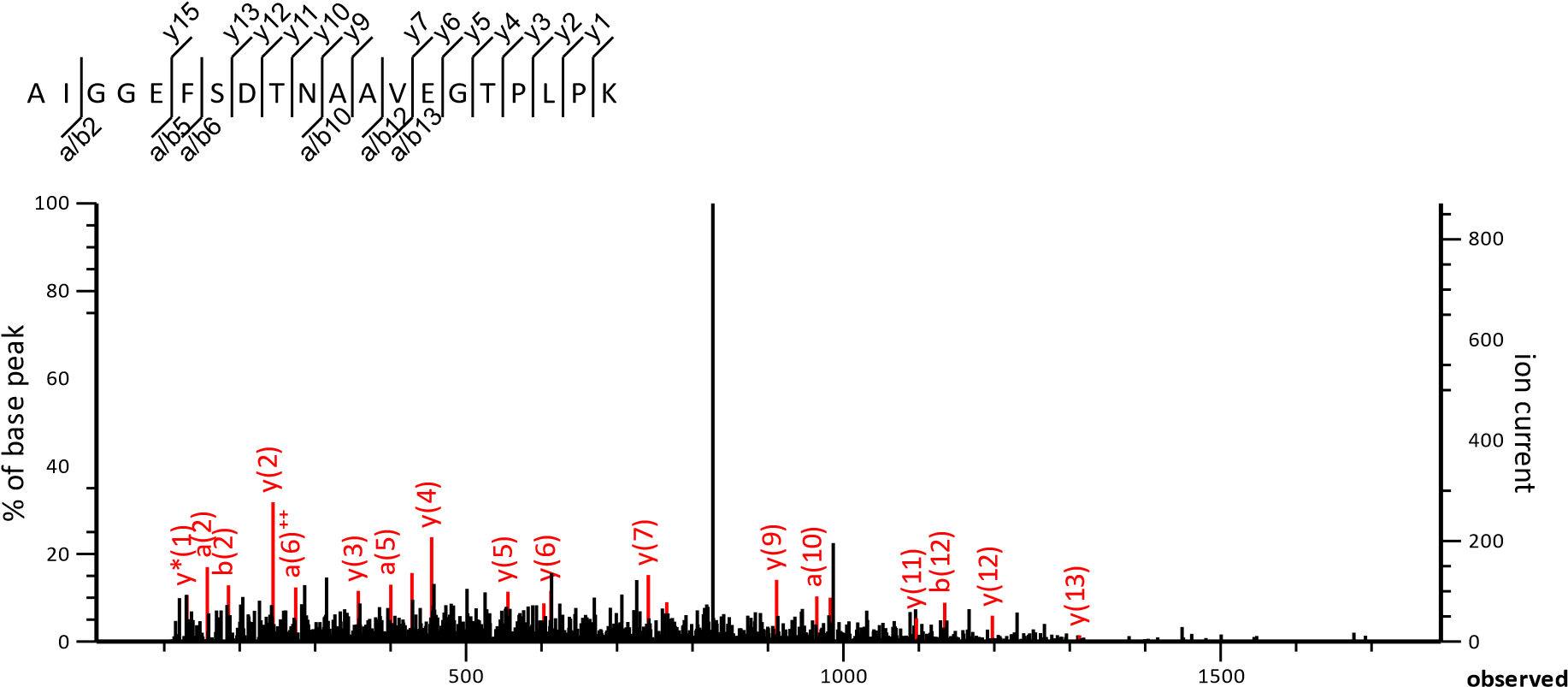

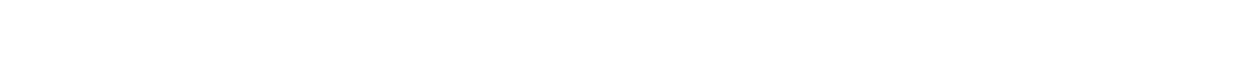

T CC1

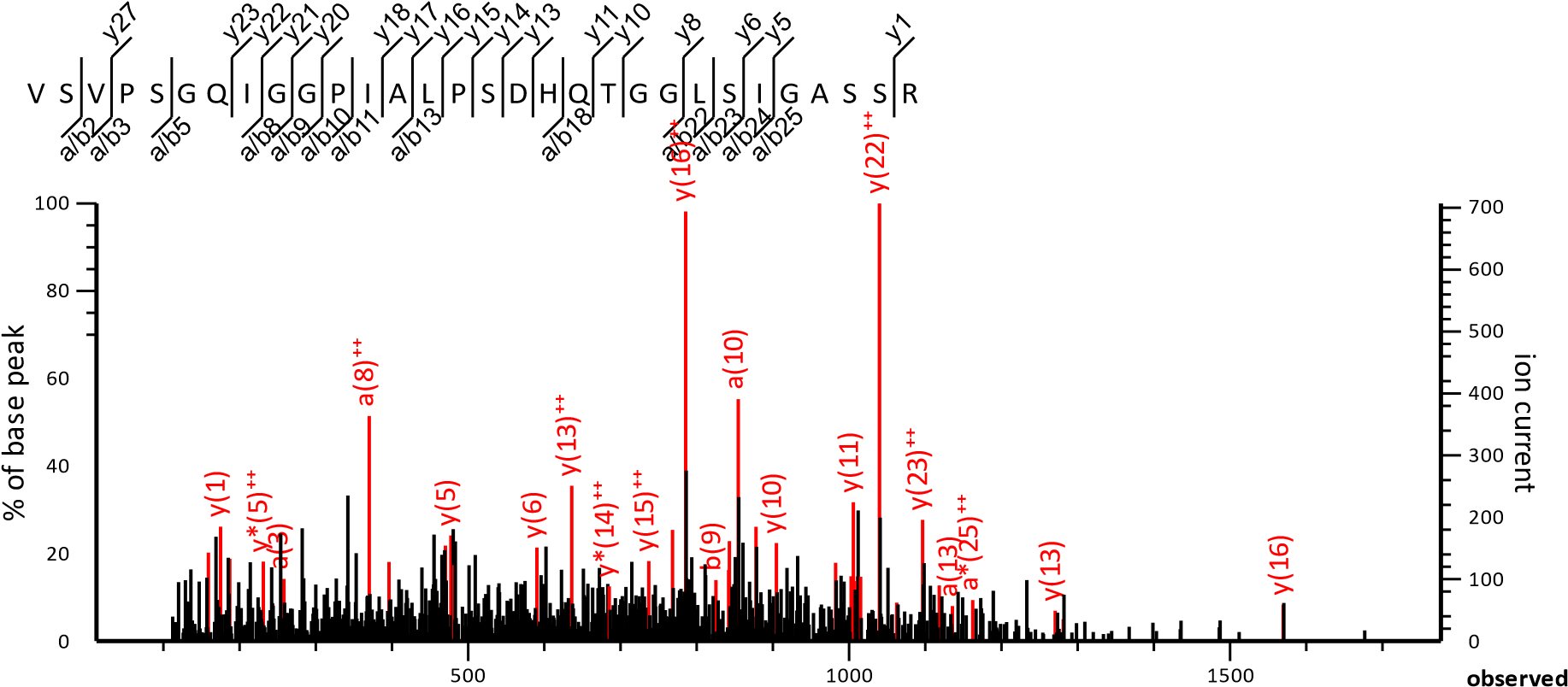

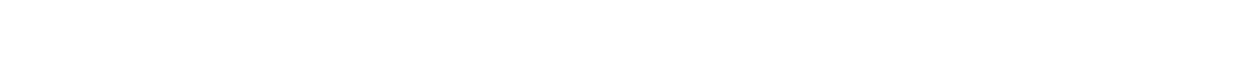

UB 1

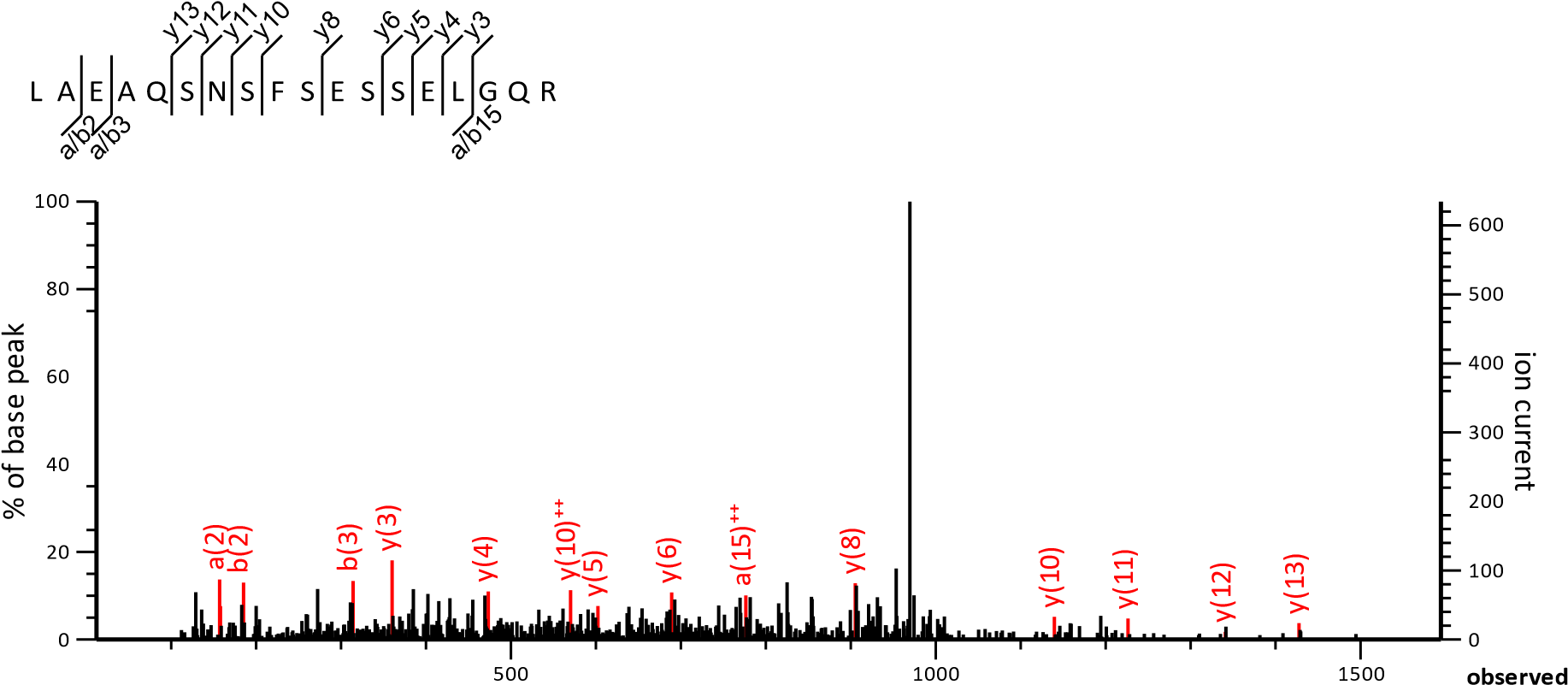

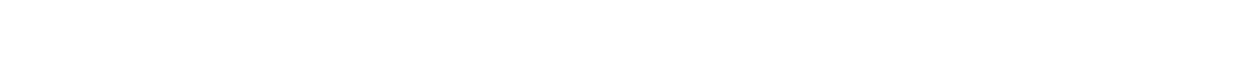

ZM T

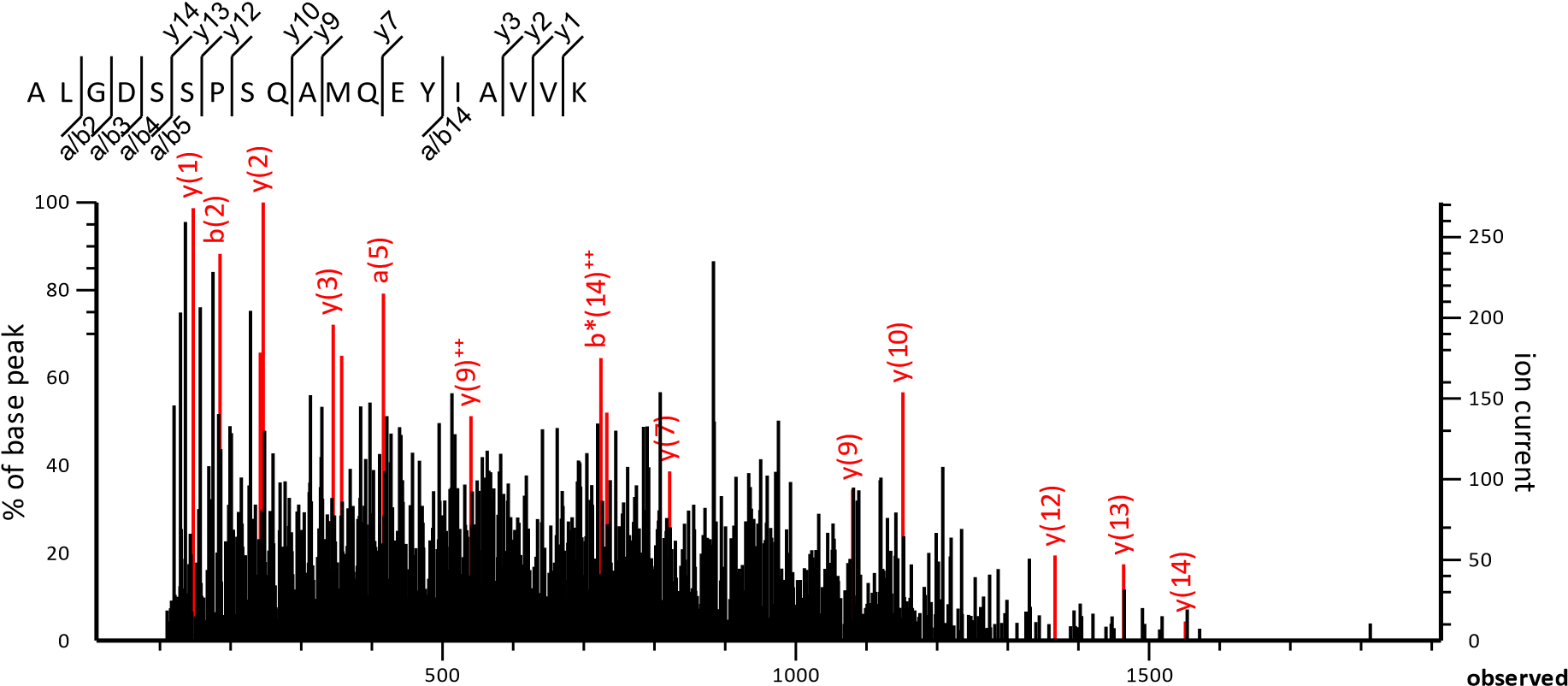

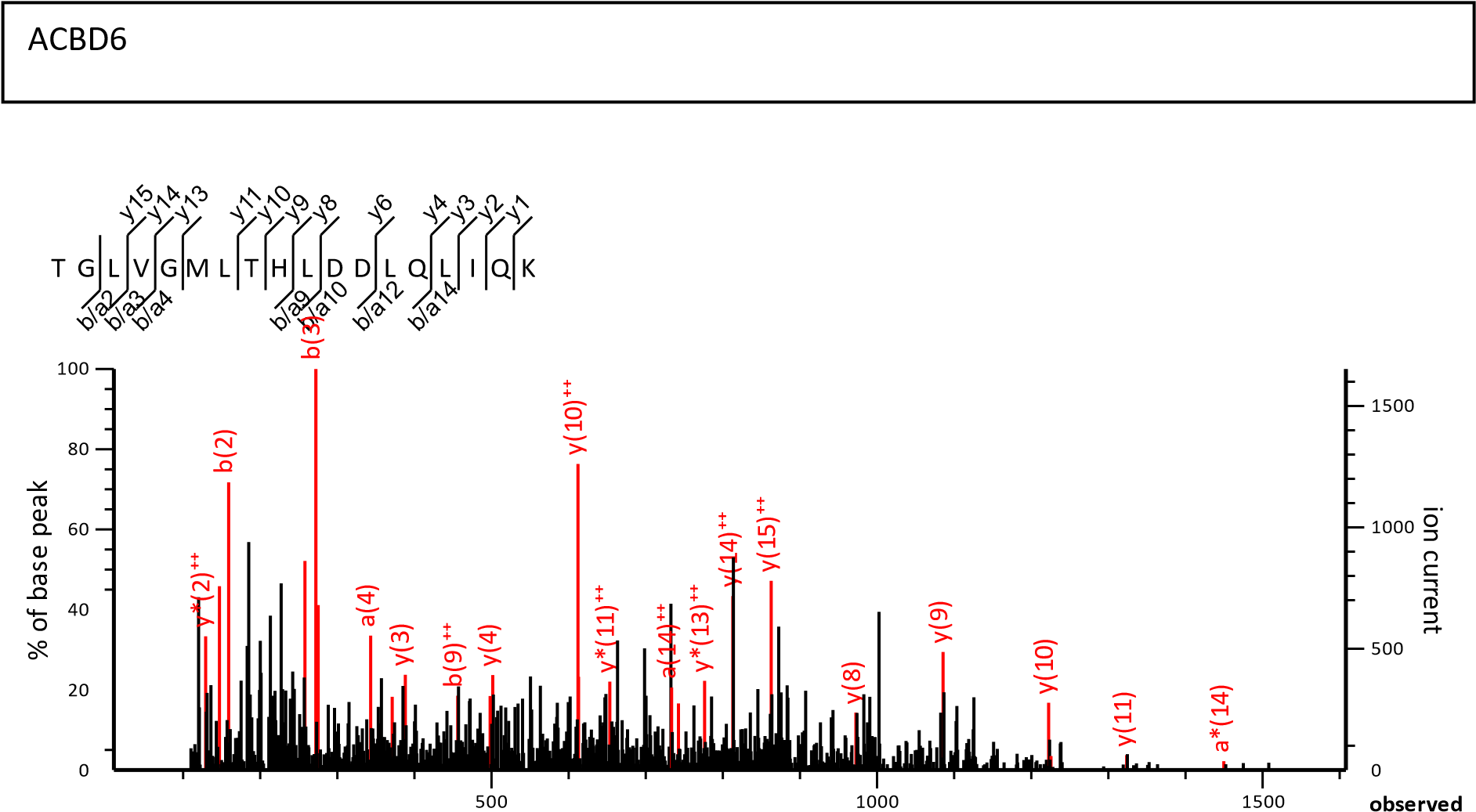

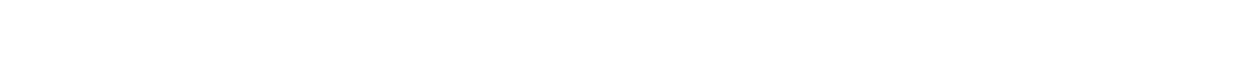

B

Fig S8. Annotated spectra for proteins identified with single peptide (Mascot search engine (version 2.7.0.1))
